# Circuit dynamics of approach-avoidance conflict in humans

**DOI:** 10.1101/2024.12.31.630927

**Authors:** Brooke R. Staveland, Julia Oberschulte, Barbara Berger, Tamas Minarik, Olivia Kim-McManus, Jon T. Willie, Peter Brunner, Mohammad Dastjerdi, Jack J. Lin, Ming Hsu, Robert T. Knight

## Abstract

Debilitating anxiety is pervasive in the modern world. Choices to approach or avoid are common in everyday life and excessive avoidance is a cardinal feature of anxiety disorders. Here, we used intracranial EEG to define a distributed prefrontal-limbic circuit supporting approach and avoidance. Presurgical epilepsy patients (n=20) performed a continuous-choice, approach-avoidance conflict decision-making task inspired by the arcade game Pac-Man, where patients trade-off harvesting rewards against potential losses from attack by the ghost. As patients approached increasing rewards and threats, we found evidence of a limbic circuit mediated by increased theta power in the hippocampus, amygdala, orbitofrontal cortex (OFC) and anterior cingulate cortex (ACC), that drops rapidly during avoidance. Theta band connectivity within this circuit and with the lateral prefrontal cortex increases during approach and falls during avoidance, and amygdala and lateral frontal activity granger-caused the theta oscillations in both the OFC and ACC. Importantly, the degree of network connectivity predicted how long patients approach, with enhanced network synchronicity extending approach times. Finally, when threat is imminent, the system dynamically switches to a sustained increase in high-frequency activity (70-150Hz) in the middle frontal gyrus (MFG), tracking the degree of threat. The results provide evidence for a distributed prefrontal-limbic circuit, mediated by theta oscillations and high frequency activity, underlying approach-avoidance conflict in humans.

## Introduction

Approach-avoidance conflict describes the daily situations where a single choice entails potential gains and losses simultaneously, which induces anxiety in everyday life (Dymond and Roche, 2009; Kirlic et al., 2017). Approach-avoidance decisions are dysregulated in generalized anxiety disorder (GAD), post-traumatic stress disorder (PTSD), and agoraphobia, where excessive avoidance of a potential aversive cue comes at the cost of reward (Ball and Gunaydin, 2022; Kirlic et al., 2017; Salters-Pedneault et al., 2004). For example, consider the decision to approach a prominent scientist in your field to offer a potential collaboration. The same action could result in the positive outcome of increasing the scientific output of your project, but could also result in social rejection and professional embarrassment. While anxiety in this context is normative, when avoidance becomes excessive (never speaking to possible collaborators nor asking for scientific guidance) it becomes maladaptive and is a core feature of all anxiety disorders (American Psychiatric Association, 2022; Dymond, 2019; Dymond and Roche, 2009). Understanding the cortical and limbic neural circuits that regulate approach and avoidance is key to understanding the neural underpinnings of both normal and excessive clinical anxiety.

There is a rich history defining the modern notion of the limbic system, which has been long known to underly aspects of emotional behavior (Catani et al., 2013; Joseph, 1992; Morgane et al., 2005), including approach-avoidance (Fried, 1971; Heilman, 1997; Vogt, 2019). In 1937, Papez described brain regions, particularly the hippocampus and anterior cingulate, associated with emotional control (Papez, 1937). Papez referred to this network of regions supporting emotion as the limbic system, based on Broca’s anatomical description of the ‘limbic lobe’ (Broca, 1878). McClean added the amygdala to the limbic system in response to Kluver and Bucy’s reports of abnormal behavior in monkeys with bilateral amygdala lesions (Klüver and Bucy, 1937; MacLean, 1952). Based on the disordered emotional behavior of patient Phineas Gage (Harlow and Miller, 1869), the orbitofrontal cortex completed the modern definition of the limbic system. Given the complexity and extent of the limbic system, we elected to interrogate the role of all these regions with a focus on theta oscillations known to be involved in network communication in a host of human behaviors. We further hypothesized that the engagement of the lateral prefrontal cortex, specifically the middle frontal gyrus (MFG), in approach-avoidance would be distinct from limbic regions (Kirlic et al., 2017; Rolle et al., 2022). The MFG has been associated with cognitive control and emotion regulation, and nonhuman primates (NHPs) research has reported evidence that the prefrontal cortex is part of a cognitive circuit exerting top-down control on the limbic circuit (Amemori et al., 2024a). Here, we use intracranial EEG (iEEG) to delineate a prefrontal-limbic system engaged in approach-avoidance including dorsolateral prefrontal cortex (dlPFC), orbitofrontal cortex (OFC), amygdala, anterior cingulate cortex (ACC) and hippocampus (HC) in humans.

Elements of this network have been explored in both rodents, nonhuman primates, and humans. Approach-avoidance conflict tasks, such as the elevated plus maze or the open field test have been used to study the neural circuits underlying anxiety in rodents (Ito and Lee, 2016; Kirlic et al., 2017; Mack et al., 2023). In these tasks, there is a conflict between the positive drive to explore new areas versus the negative drive to avoid exposed areas. Using a variety of approaches, researchers have identified pairs of prefrontal-limbic regions, mediated by theta-band power, associated with avoidance behavior on these tasks (Calhoon and Tye, 2015; Jacinto et al., 2016; Padilla-Coreano et al., 2019). Researchers have found that a subset of ventral hippocampal (vHC) neurons that project to the medial PFC (mPFC) show increased theta synchrony in anxiogenic contexts (Adhikari et al., 2010b). Optogenetically stimulating these neurons at 4-12Hz increases both synchrony between the vHC and the mPFC and increases avoidance behavior in rodents (Padilla-Coreano et al., 2019). Similarly, optical activation of rodent basolateral amygdala neurons that terminate in the mPFC increases avoidance in the elevated plus maze, open field test, and other anxiogenic tasks, while inhibition of these neurons increases approach behavior (Felix-Ortiz et al., 2016). Other studies report bidirectional mPFC-amygdala circuitry, implicating the mPFC in top-down control of amygdala-mediated anxiety behaviors in rodents (Adhikari et al., 2015; Mack et al., 2023). However, the human homologue of the rodent mPFC is still debated (Laubach et al., 2018)and the massive expansion of human prefrontal cortex limits a direct translation of rodent to human findings.

There is evidence that similar regions are engaged during approach-avoidance in humans using both non-invasive imaging and lesioned patients. Bilateral hippocampal lesioned patients showed increased approach behavior (Bach et al., 2019), and a magnetoencephalography study found that hippocampal-PFC theta synchrony increased with threat (Khemka et al., 2017). Some fMRI studies of human approach-avoidance conflict report activations in the amygdala (Abivardi et al., 2020), but others do not (Aupperle et al., 2011; Schlund et al., 2016; Talmi et al., 2009). This has led researchers to hypothesize that the amygdala responds equally to appetitive and aversive cues and may be missed in standard fMRI contrasts (Kirlic et al., 2017). Other studies using fMRI have found activations in orbitofrontal cortex, anterior cingulate and dorsolateral prefrontal cortex that correlate with task parameters like increasing threat, reward, and conflict, though there is considerable variation between different tasks (Chu et al., 2023; Kirlic et al., 2017; Schlund et al., 2016; Talmi et al., 2009; Zorowitz et al., 2019). Separately, researchers have found that midline frontal theta, as measured via EEG, increases with approach-avoidance conflict and might be related to the midline frontal theta elevations in other studies requiring cognitive control (Lange et al., 2022).Together, this work suggests that both prefrontal and limbic regions are engaged in approach-avoidance behavior. However, the real-time circuit dynamics of approach-avoidance has yet to be characterized in humans.

Here we test for the existence of such a prefrontal-limbic network underlying approach-avoidance by examining 1) task-dependent modulations in theta power in middle frontal gyrus and limbic regions including the hippocampus, amygdala, OFC and anterior cingulate; 2) elevated theta band coherence across these regions; and 3) dynamic changes in this circuit in response to increases in threat and reward. To test these hypotheses, we collected intracranial electrode recordings across this circuit in twenty epileptic patients during presurgical assessment while they performed a continuous-choice, approach-avoidance conflict task based on the arcade game Pac-Man.

## Results

### Behavioral Results

We designed an approach-avoidance conflict task based on the arcade game Pac-Man (See Figure 1a). At trial onset, Pac-Man is placed along a single corridor that runs horizontally across the screen. The player makes a button press (either the left or right arrow key) to begin moving along the corridor to the left or the right. They can either choose to move towards the *end* of the corridor where they can exit to the next trial, or they can move towards the *center* of the corridor where the patient encounters both potential gains (eating “dots”, resulting in points) and potential losses (ghost attack, resulting in loss of a life of Pac-Man). Patients could choose to eat up to 5 dots, which varied in either small (10 points) or large (20 point) sizes, on each trial, but they could also choose to turn back towards the end of the corridor at any time, thereby exiting the trial. Being chased or being caught by the ghost are determined by the player’s distance to the ghost. The participants were instructed to balance the risks of being caught by the ghost and the rewards of getting as many dots as possible to achieve the highest score. If the ghost consumes Pac-Man, all points acquired in that trial are lost. Direction, reward and location of the ghost and Pac-Man are counterbalanced across trials. 20% of trials are conflict-free, where the player is free to collect the 5 dots without threat of the ghost.

**Fig. 1.**
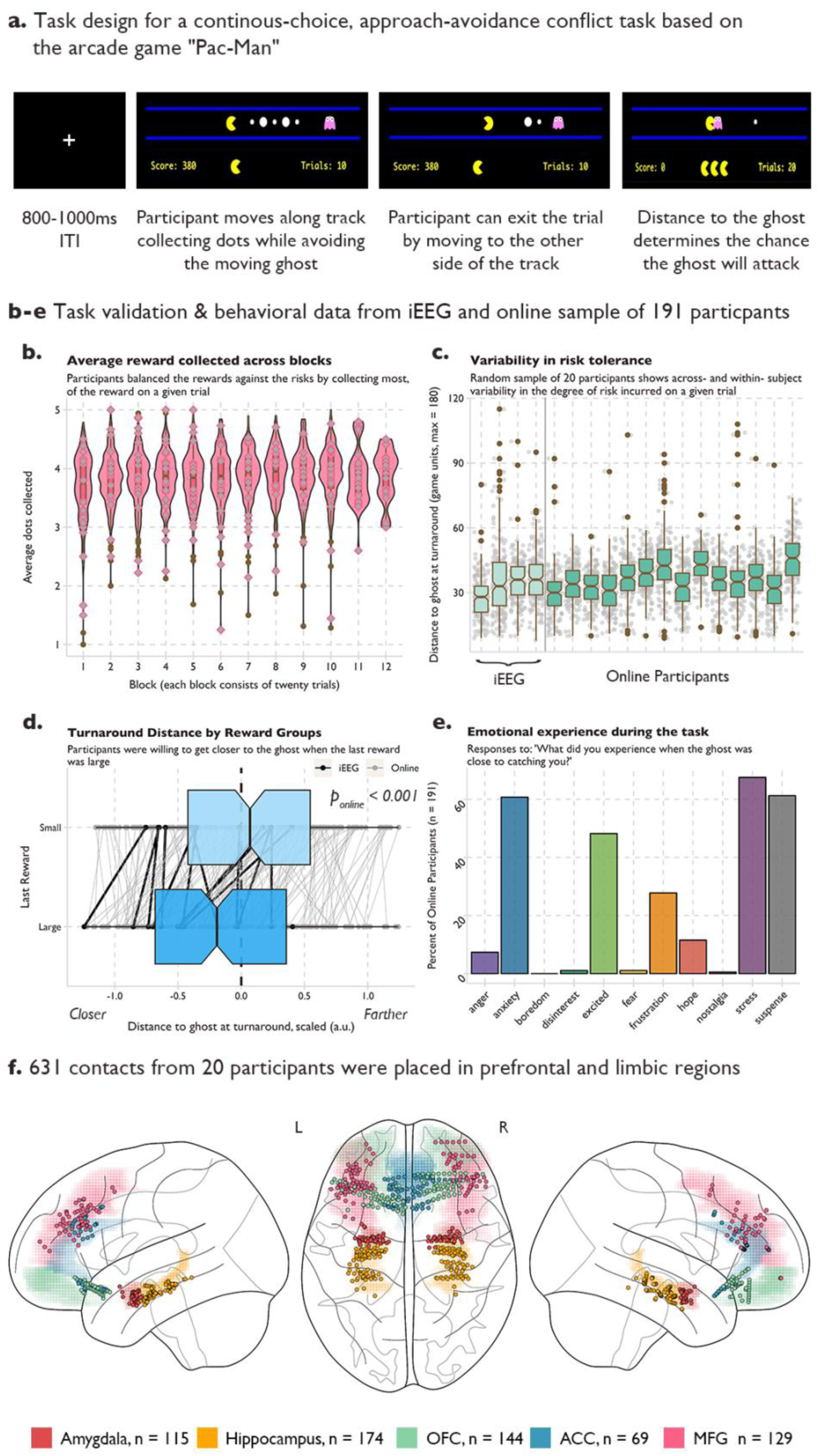
**a** Task design for a real-time, approach-avoidance conflict task. Pac-Man was placed along a horizontal corridor, with a ghost moving in a set path back and forth at the opposite end of the corridor. Five rewards of varying sizes were placed between Pac-Man and the ghost. Participants used the left and right arrow keys to move closer to the ghost to collect reward (approach), or to move back to the end of the corridor where participants could exit the trial (avoid). **b-e** Normative behavior on the task in iEEG patients and an online sample of 191 participants. **b** Average reward collected across blocks. Violin and box plots of the average number of rewards (dots) collected on each trial across 12 blocks of 20 trials. Grey diamonds indicate the performance of each intracranial participant. Variability in risk tolerance. **c** Box plots for 20 example participants and 4 intracranial participants of the average distance to the ghost at the final decision to avoid across all trials. Grey dots indicate the turnaround distance for each trial. **d** Turnaround distance by reward group. Rewards could be large (worth 20 points) or small (worth 10 points). On trials where the last dot was large, participants moved closer to the ghost compared to trials where the last reward was small. Black dots indicate average distance for each intracranial patient, grey dots indicate the average for each online participant, and the lines link the behavior within participants. **e** Emotional experience of the task in the online sample. Y-axis is the percent of participants who reported experiencing the emotion on the X-axis. Participants were allowed to select more than one emotional experience or write their own. **f** Electrode placement. 631 electrodes were placed across the prefrontal and limbic regions in 20 intracranial participants. Colored shading indicates region and dots indicate electrode.

We validated that the task was inducing approach-avoidance trade-offs by assessing the behavior in an online sample of 191 participants recruited from the online platform Prolific (see Methods for details). Across the twelve blocks of twenty trials each, participants collected an average of 3.83 +/ 0.34 dots per trial, demonstrating that they often left some potential reward in the trial (max 5 dots) to maintain a greater distance from the ghost (See Figure 1b). Participants also demonstrated individual variance in their risk tolerance as the variability in this distance within a participant was smaller than the variability across participants (See Figure 1c), indicating that participants varied in how close they were willing to get to the ghost to receive more reward. We also hypothesized that participants would make risk/reward trade-offs tolerating more risk to get large rewards. As expected, when the last reward in the corridor was large, participants were willing to move closer to the ghost to collect the large reward (See Figure 1d). Since approach-avoidance conflict is often used as a behavioral model of anxiety, we asked participants to report their emotions during the trial. Participants were given a set of emotions (anger, anxiety, boredom, disinterest, excitement, frustration, hope, stress, and suspense) as well as an option to write their own. Participants could report multiple emotions. A majority of participants reported experiencing anxiety, stress, and suspense during the task, supporting face-validity of the task (Fig 1e).

Twenty patients with intracranial electrodes in the amygdala, hippocampus, OFC, ACC, and MFG (Fig 1e; see methods and supplementary data for patient details) performed the task. Behavioral performance across the twenty patients was comparable to the performance in our online sample (See Figure 1 b-e). Electrodes near seizure foci were removed, and epochs where seizure activity spread beyond the local seizure onset zone were excluded. This left a total of 115 amygdala contacts, 174 hippocampal contacts, 144 OFC contacts, 69 anterior cingulate contacts, and 129 MFG contacts that were included in our analyses. While some patients did not have contacts in all five regions, there was always at least 12 patients with coverage in each possible *pair* of our five regions, allowing us to probe the neural dynamics across all sets of regions (See Supplemental Figure 1 for individual patient electrode coverage).

### Theta power is locally modulated by approach-avoidance decisions in limbic regions

We hypothesized there would be task-dependent theta power modulations within our regions of interest. In each region, we calculated time-frequency plots time-locked to the patients’ choice to stop approaching and turn back to exit the trial. We observed a similar pattern in the hippocampus, amygdala, OFC, and ACC, where low-frequency power between 3-8Hz, is elevated during the approach period of the trial but drops after the decision to avoid and exit the trial (See Figure 2, Panel A).

**Fig. 2.**
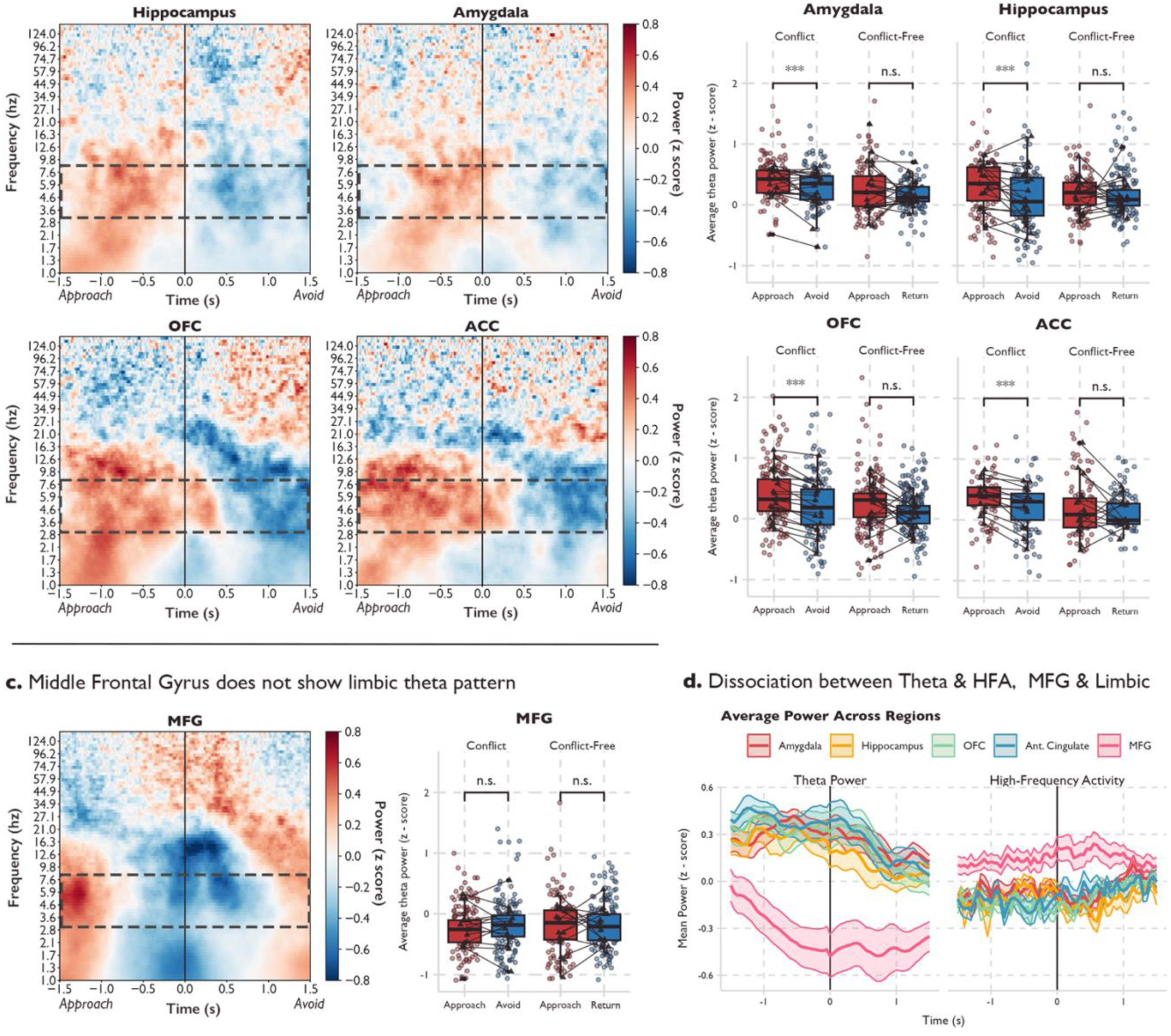
**a** Time-frequency analysis of neural activity time-locked to the final choice to stop approaching and begin avoiding across all electrodes in the limbic regions (hippocampus, amygdala, OFC and ACC). Red (blue) indicates increases (decreases) in power. Power was log-transformed and z-scored to the mean power across the 3-second time window. Dotted box indicates the theta band (3-8Hz). **b** Average theta power in the 1500ms before and after the choice to avoid for each limbic region in conflict-trials (ghost present) or conflict-free trials (ghost absent). Dots indicate average theta power in electrodes across trials, triangles indicate average theta power across electrodes within a participant. Significance was assessed using linear-mixed effects models with random effects of participant and electrode. Stars indicate that the 95% credible interval did not include 0. **c** Time-frequency analysis and theta-power analysis for the MFG. **d** Time-course of theta power (left subplot) and high-frequency activity (HFA, right-subplot) across approach and avoidance windows. Power data was extracted from the TFR and z-scored using the average power in the ITI. Color indicates region, shading is the standard error of the mean power across patients.

We next averaged theta power during the final 1500 milliseconds of approach and first 1500 milliseconds of avoidance in each electrode and patient. We estimated a Bayesian mixed effects model of theta power using their approach/avoidance decisions with hierarchically grouped random effects of patient and electrode (See Methods for Details). We found that theta power was elevated during approach compared to avoidance in trials where there was a threat within the hippocampus (See Figure 2, Panel a; Conflict-Trial Estimate_approach>avoidance_: 0.20, 95% CI [0.06-0.35]), amygdala (Conflict-Trial Estimate_approach>avoidance_: 0.14, 95% CI [0.03-0.25]), OFC (Conflict-Trial Estimate_approach>avoidance_: 0.18, 95% CI [0.11-0.26]) and ACC (Conflict-Trial Estimate_approach>avoidance_: 0.19, 95% CI [0.08-0.29]). In the subset of control trials that were conflict-free, i.e. where patients collected reward without threat of the ghost, we did not find a significant difference in theta power between the approach and avoidance windows (Hippocampus Estimate_approach>return_: 0.03, 95% CI [-0.16-0.23]; Amygdala Estimate_approach>return_: 0.10, 95% CI [-0.05-0.23]; OFC Estimate_approach>return_: 0.13, 95% CI [-.06-0.32]; ACC Estimate_approach>return_: 0.09, 95% CI [-0.16-0.34]), indicating that the effects were not likely driven by purely spatial or reward-only behavior (See Supplemental Figure 2 for TFRs of the conflict-free trials).

We then tested for theta power modulations in the MFG, implicated in previous fMRI and nonhuman-primate studies of approach-avoidance but not considered to be part of the limbic system (Amemori et al., 2024b; Chu et al., 2023; Rolle et al., 2022). We did not find evidence for increased theta power in the approach period compared to the avoidance period in the MFG (Conflict-Trial Estimate_approach>avoidance_: 0.03, 95% CI [-0.10-0.16]; Conflict-Free Estimate_approach>return_: -0.12, 95% CI [-0.32-0.10]). We next tested for the existence of theta oscillations in our MFG contacts, to understand if the absence of theta power changes between approach and avoidance windows was due to a lack of theta oscillations in this region, or due to inconsistency across patients or power stability across the trial. We used the FOOOF (Fitting Oscillations and One-Over-F) algorithm—which decomposes the power spectrum into its aperiodic (1/f) and periodic components— to these contacts and found evidence of theta oscillations in the MFG in each patient in at least one contact (Donoghue et al., 2020). Therefore, while there are theta oscillations in the MFG in our task, the dynamics of the theta oscillations did not differ between approach and avoidance periods and they were inconsistent across contacts and patients (see Supplemental Figure 3). Instead, the MFG was characterized by an elevation of high-frequency activity (70-150 Hz) around the time of choice not apparent in other regions (See Figure 2c).

### Limbic regions are connected via theta band activity during approach-avoidance decision-making

We hypothesized that the modulations of local theta power were accompanied by increased connectivity between these regions. We calculated the imaginary part of coherence, which removes the interactions due to potentially spurious volume conduction effects (Bastos and Schoffelen, 2016), between theta activity in pairs of electrodes from different regions. We compared the true imaginary coherence to a shuffled distribution where trial labels in one region were shuffled 1000 times. Each time point was compared to the shuffled distribution and then FDR-corrected for the number of time points. Pairs of electrodes were marked as having significant theta-band coherence if there was at least a time window of 100ms of significant elevated coherence compared to the null distribution (See Figure 3a). We also calculated two alternative coherence measures, pairwise phase consistency and the debiased estimator of the weighted phase lag index that confirmed our results were robust across different estimates of coherence.

**Fig. 3.**
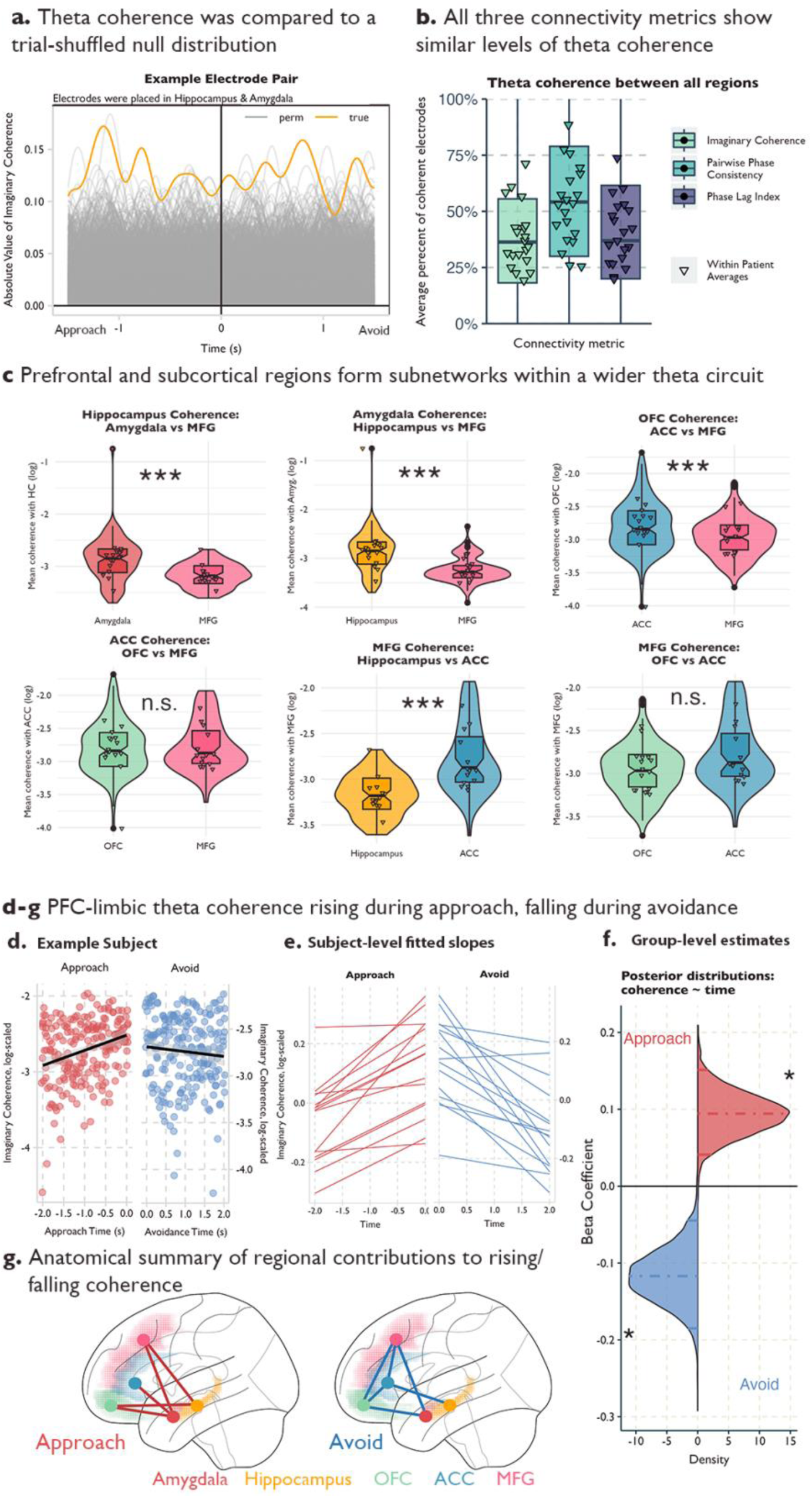
**a** Example coherence in an electrode pair from a participant with contacts in the hippocampus and the amygdala. Yellow line indicates the true absolute value of imaginary coherence across the approach and avoidance periods. Grey lines indicate the absolute value of imaginary coherence from the permuted sample, where trial labels for the amygdala contact were shuffled 1000 times. **b** Average percentage of electrode pairs with elevated theta coherence out of all possible pairs for the three connectivity metrics. Triangles denote patient-level averages, color denotes connectivity metric. **c** Violin and boxplots for comparing theta coherence strength between two partner regions within each region of interest. Each facet is a different region of interest, and we have highlighted two partner regions to compare on the x-axis (See Supplemental Figure 4 for all region comparisons). Triangles indicate patient averages, *** indicates the 95% confidence interval of the estimated difference in connectivity strength between the two regions did not include 0. **d-g** Increase (decrease) in theta coherence during the approach (avoidance) window. **d** Example patient. Y-axis is the log of the absolute value of imaginary coherence averaged across contacts and region pairs in the approach (red) avoidance (blue) period. X-axis is time in seconds and each period is fit with a regression line. Shading indicates 95% confidence intervals. **e** Patient-level fitted slopes representing the effect of time on theta coherence in the approach period (red, left) or avoidance period (blue, right). **f** Posterior distributions for the effect of time on theta coherence in the approach period (red) or avoidance period (blue). **g** Regional differences in the effect of time on theta coherence in the approach (left) or avoidance (right) windows. Red (blue) lines indicate that the connected regions significantly drove the effect in approach (avoidance) time windows. Colored shading indicates region label.

We found more electrode pairs demonstrated coherence within the theta band than would be predicted by chance across each of the limbic regions and with the middle frontal gyrus (See Figure 3b). We calculated the percentage of electrode pairs that showed elevated coherence in the theta band and averaged this percentage across our 20 patients. In all regions, on average, at least one-third of the pairs showed elevated coherence (See Supplementary Table 1), which is higher than the percentage expected by chance. This was similar across all three measures of connectivity.

Having established the existence of theta-band coherence across our regions of interest, we then tested that the consistent limbic theta *power* modulations were indicative of elevated theta *coherence* between limbic regions, and would show increased connectivity with each other compared to the MFG. To test this, we ran a Bayesian mixed effects model within each region to predict theta coherence, using a four-level factor representing the remaining regions as the predictor. This analysis can be thought of as a frequentist ANOVA; however, because the entire model is estimated at once, pairwise differences in coherence strength can be assessed directly without the need for post-hoc tests. We included patients and electrodes as random effects. To avoid inflating our estimates, we tested these models across all electrode pairs, even those that did not show elevated theta coherence in our previous analysis.

As predicted, we found that the amygdala exhibited higher theta coherence with the hippocampus compared to the MFG, with an estimated connectivity strength ratio of 1.49 (95% credible interval [1.29, 1.75], See Figure 3c). This indicates that amygdala-hippocampus coherence was ∼1.5x stronger than amygdala-MFG coherence during the approach. Similarly, we found that amygdala-OFC coherence and amygdala-ACC coherence were both stronger than amygdala-MFG coherence, with a connectivity strength ratio of 1.22 (95% credible interval [1.07, 1.36]; see Supplemental Figure 4) and 1.14 (95% credible interval [1.03, 1.28]; see Supplemental Figure 4), respectively. We found similar results in the hippocampus (See Figure 3c; Supplemental Figure 4; Supplemental Table 2); hippocampus coherence with the amygdala was 1.41 (95% credible interval [1.23, 1.65]) times greater than its theta coherence with the MFG, and 1.14 (95% credible interval [1.03, 1.27]) times greater with the OFC compared to the MFG. These results provide some evidence that the subcortical limbic regions (e.g., amygdala and hippocampus) are more strongly connected to other limbic regions via theta compared to the MFG (See Supplemental Figure 4 for all regional comparisons using Imaginary Coherence; see Supplemental Figure 5 for results using Pairwise Phase Consistency and Phase Lag Index; see Supplemental Table 2 for full model results across the three connectivity metrics; see Supplemental Table 3 for posterior predictive checks across connectivity models).

When we turn our attention to the prefrontal limbic regions (e.g., OFC and ACC) a more complex network structure develops. In the ACC, we see stronger theta coherence with the MFG compared to the amygdala and hippocampus, where ACC-amygdala coherence was only 0.76 (95% credible interval [0.65, 0.88]) times the ACC-MFG coherence and ACC-hippocampus was only 0.77 (95% credible interval [0.65, 0.90]) times ACC-MFG coherence (See Figure 3c, Supplemental Figure 4 and Supplementary Table 2 for details). Similarly, the ACC showed preferential connectivity with the OFC compared to the subcortical limbic regions (See Figure 3c, Supplemental Figure 4 and Supplementary Table 2 for details), meaning that, while the ACC shared similar theta power modulations as the subcortical limbic regions, the ACC was preferentially connected to other prefrontal regions compared to the two subcortical regions.

Furthermore, we found that the MFG also had preferential connectivity with the other prefrontal regions (OFC, ACC) compared to subcortical regions (See Figure 3c, Supplemental Figure 4 and Supplementary Table 2 for details), matching the profile we found in the ACC. While both the ACC and MFG and the hippocampus and amygdala shared similar connectivity profiles during approach, the OFC had a more distributed connectivity. There were no significant differences between its connectivity with subcortical regions compared to the MFG, though there was a preference for ACC connectivity compared to these other regions (See Figure 3c, Supplementary Figure 4, Supplementary Figure 5, Supplementary Table 2-3 for details).

### Theta coherence increases over the approach period and falls during avoidance

Next, we examined dynamic task-related changes in this circuit. Rodent models report that theta synchrony between the hippocampus and mPFC increases in anxiogenic versus safe contexts (Adhikari et al., 2010b). We hypothesized that theta synchrony across the prefrontal-limbic network would increase across the approach period as patients increased both the level of reward, threat and conflict.

We tested this by using time to predict theta coherence in the two seconds preceding the choice to avoid. We averaged the connectivity values by region pair (e.g., OFC-ACC, Amyg.-Hipp., etc.) within patients and within 100ms time bins. We again used Bayesian mixed-effects models with a random effect of patient and electrode pair and limited electrode pairs to only those that showed significant levels of theta coherence in either the approach or avoidance periods to ensure we were capturing fluctuations in true coherence as opposed to random noise. As hypothesized, we find the average connectivity values are increasing during the time the patient approaches (Estimate: 0.09, 95% CI [0.05-0.14]; See Figure 3d-g). While local theta power might potentially affect connectivity results this is unlikely the case because theta power – even though initially elevated – drops (See Figure 2c) during the time interval of increased connectivity. We confirmed this result using two other connectivity metrics (Pairwise Phase Consistency, Phase Lag Index) and across four different time windows ranging from 1.5-2 seconds (Supplementary Table 4). We next included an interaction for region pairs to test for specific region pairs that were driving this effect. We found that the subcortical-prefrontal pairs including amygdala-MFG, amygdala-ACC, amygdala-OFC, hippocampus-MFG, and hippocampus-OFC demonstrated strong time-coherence modulations (See Figure 3g). Next, we tested if theta coherence would fall, remain constant, or continue to rise during the avoidance period. We hypothesized the coherence would fall when retreating from the ghost, indicating that theta coherence across the circuit is highest during the portion of the trial with the highest levels of risk and conflict. As hypothesized, we found that theta coherence drops after the patients began avoiding and moved to exit the trial (Estimate: -0.12, 95% CI [-0.18 - -0.04]), and this effect was driven by both subcortical-prefrontal pairs (amygdala-MFG, amygdala-OFC, hippocampus-ACC), and also prefrontal-prefrontal pairs (MFG-ACC, MFG-OFC, OFC-ACC; See Figure 3d-f; Supplementary Table 4). As before, we found that the drop in theta coherence during avoidance was consistent across both alternate connectivity measures and multiple timepoints (See Supplementary Table 4).

### Amygdala and lateral prefrontal activity drive theta oscillations in the OFC and ACC

We next assessed the directionality of the prefrontal-limbic circuit using two independent methods, State-Space spectral granger causality and cross-correlation analysis (Adhikari et al., 2010a; Barnett and Seth, 2015). We calculated the net spectral Granger causality between all electrode pairs with significant levels of theta-band coherence in the last 1500ms of the approach period. As there may be bidirectional information flow, we identified which signals dominated the information flow by calculating the difference between the Granger values calculated in both directions. We next subtracted the spectral Granger values calculated from the reversed time-series, as noise can inflate connectivity estimates if the signal-to-noise-ratio SNR increases or decreases over time (Haufe et al., 2013). This final net Granger score was compared to a permuted distribution where trial labels from one electrode were shuffled 200 times. By comparing the absolute value of the true net Granger score to the permuted null distribution, we calculated permuted p values and then FDR-corrected these values for the number of timepoints. If more than 50 timepoints (500ms) were significant at p < .05, then that electrode pair was included in our subsequent analyses.

When net Granger scores were higher than chance, we tested for a bias in directionality between our regions. We turned to our linear mixed-effects models to account for shared variance within electrodes and patients. Specifically, we modeled the net Granger scores during the approach window using an intercept-only model and random effects of electrode and patient. From these models, we report the Probability Positive (P+), which represents the proportion of posterior samples where the effect is greater than zero, as a measure of directional certainty, and we consider P+ > 0.95 as strong evidence for directionality. Since we did not have strong *a priori* hypotheses about the directionality in this circuit, we also validated these analyses by conducting a cross-correlation analysis on the theta-band passed signals for each pair of cohering electrodes. We used similar linear mixed effects models to assess the P+ of the lead and lag times resulting from the cross-correlation analysis. Results were only interpreted if they resulted in significant P+ values using both independent methods, which relied on different calculations of theta power (Morlet Wavelets vs Bandpass filtering) and slightly different time windows (the cross-correlation analysis excluded timepoints where the patient was still at the beginning of the trial, while the Granger analysis included these timepoints). Details of both analyses can be found in the Methods.

While we found evidence of bidirectional information flow in many individual pairs of electrodes, we found four region pairs with consistent directionality across our patients. First, we found that theta oscillations in the Amygdala drove theta in both the OFC (P_Granger_+ = 0.97; P_CCF_+ = 0.96) and ACC (P_Granger_+ = 0.98; P_CCF_+ = 0.96). Second, we found that the Middle Frontal Gyrus also led the theta oscillations in the OFC (P_Granger_+ = 0.95; P_CCF_+ = 0.98) and ACC (P_Granger_+ = 0.99; P_CCF_+ = 0.99), meaning that both the OFC and ACC were being driven by subcortical and lateral prefrontal contributions. We did not find consistent evidence of directionality between any of the other region pairs (Amyg -> HC: P_Granger_+ = 0.92; P_CCF_+ = 0.70; Amyg -> MFG: P_Granger_+ = 0.64; P_CCF_+ = 0.81; HC -> OFC: P_Granger_+ = 0.93; P_CCF_+ = 0.53; HC -> ACC: P_Granger_+ = 0.65; P_CCF_+ = 0.05; HC -> MFG: P_Granger_+ = 0.58; P_CCF_+ = 0.16; OFC -> ACC: P_Granger_+ = 0.42; P_CCF_+ = 0.78).

### Pairwise synchrony during approach behavior predicts choice to avoidance

We hypothesized that theta synchrony across the prefrontal-limbic circuit during approach would correlate with the time spent approaching given rodent data suggesting a role for theta in approach-avoidance (Adhikari et al., 2010b; Padilla-Coreano et al., 2019). Coherence measures were not reliable at the trial level, and we instead calculated the Pearson correlation between the amplitude envelopes of the theta signals in each electrode pair during the approach period, which is another common method for assessing functional connectivity between regions (Mercier et al., 2022). We tested this using Bayesian mixed effects models with random effects of patient and electrode in the set of electrode pairs that showed significantly high theta coherence. As hypothesized, we found that coordinated theta activity within numerous region pairs in this circuit correlated with approach times (See Figure 4 a-b). In particular, theta synchrony between the OFC and all other region pairs correlated with longer approach times.

**Fig. 4.**
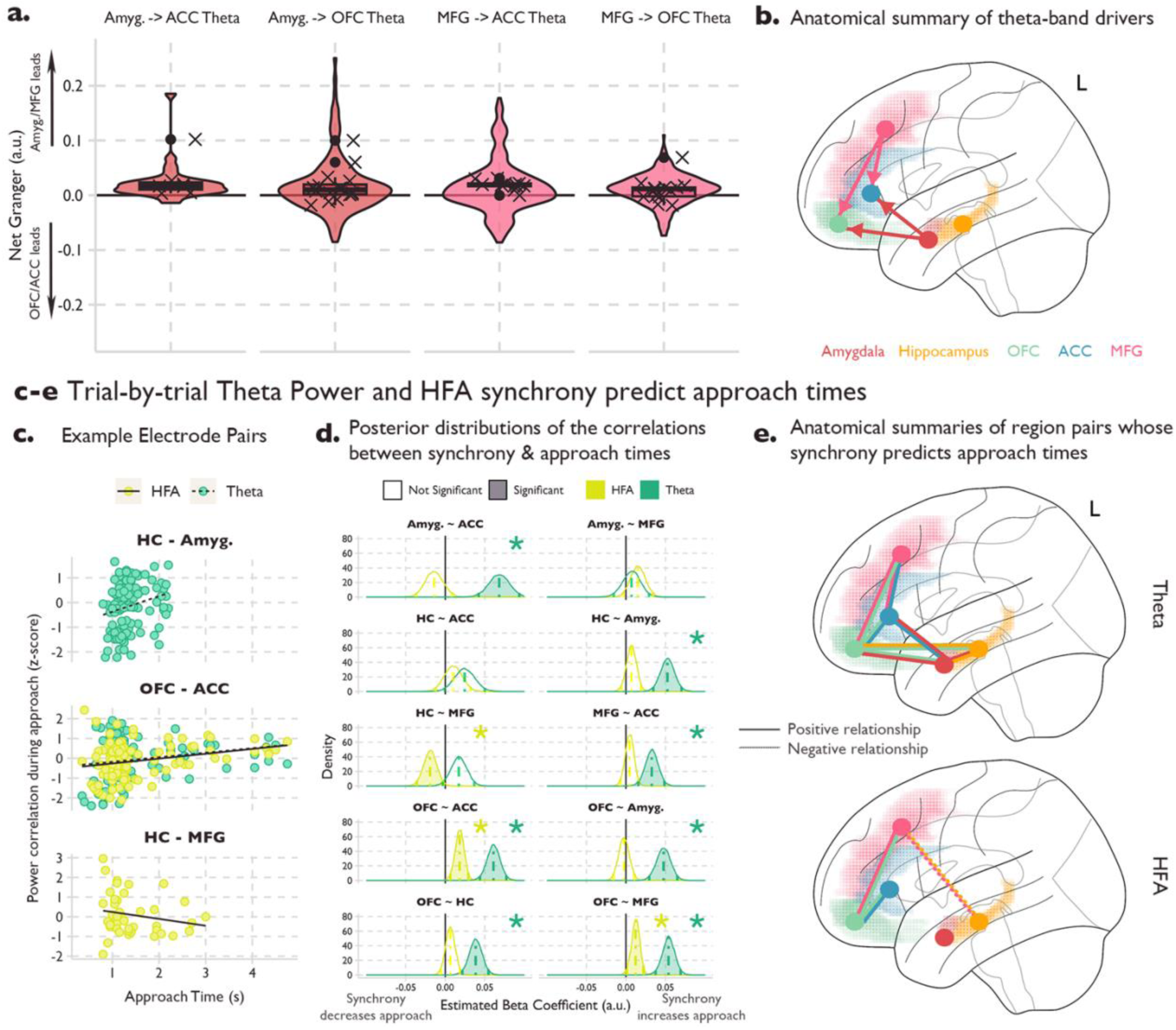
**a-b** Results from the spectral Granger causality (GC) analysis between 4 sets of regions. Y-axis shows net GC and each facet shows a region pair. Violin and boxplots show the average granger values across the approach window. Crosses show the patient averages. Positive values indicate that either the Amygdala or MFG led, while negative values indicate that either the OFC or ACC was the leading region. **b** Anatomical summary of the directional connectivity analysis. Color corresponds to region, and arrows denote the directionality. **c-e** Trial-by-trial theta and HFA synchrony predicts approach times. **c** Association between approach times and theta/HFA synchrony in example pairs of electrodes. X-axis is the untransformed time that a participant spent approaching on a given trial. Y-axis is the z-scored correlation of theta power during the approach period between the two contacts. Each facet is from a different example pair, and color indicates frequency band. **d** Posterior distributions of the estimated correlation between approach times and either theta power (green) or HFA (yellow) synchrony. Linear mixed-effects models were fit using MCMC sampling to estimate the association between approach times and theta synchrony. Colored, dotted lines indicate the mean of the posterior distribution, and solid color lines indicate the 95% credible intervals. Both stars and transparency indicate if the credible intervals include 0. **e** Anatomical summary of the results presented in d. Lines indicate that the connected regions predicted approach times. Solid lines indicate a positive relationship with approach time, dotted lines indicate a negative relationship. Top (bottom) brain shows results for theta-band (HFA) results.

These data provide strong evidence that a distributed prefrontal-limbic theta circuit is recruited during approach-avoidance conflict, with increased theta coherence associated with longer approach times. We hypothesized that theta power was providing a mechanism for network communication and next tested if local neuronal firing (Leszczyński et al., 2020; Rich and Wallis, 2017), as approximated by high-frequency activity (HFA), also correlated with approach times. We used the same high theta coherence electrodes and calculated the Pearson correlation of the high-frequency activity during approach between the electrodes. Here, we found a more restricted set of region pairs that were predictive of approach time.

Specifically, we found that OFC-MFG and OFC-ACC high-frequency synchrony correlated with longer approach times. We also found a negative correlation between hippocampus-MFG high-frequency synchrony and approach times indicating that the more similar the high-frequency activity in the hippocampus and MFG, the more likely the patient was to turn and avoid attack earlier (See Figure 4 c-d).

### Network circuit reorganization after decision to avoid

We have found evidence of a theta-mediated circuit across cortico-limbic regions which increases in connectivity strength as reward, threat, and conflict increases. Additionally, we have found that synchrony in both the theta band and HFA in these regions predicts approach times and that the OFC connects the prefrontal regions with the hippocampus and amygdala. Next, we aimed to understand how this system changes as threat becomes more proximal. It has been proposed that separate neural circuits and corresponding behaviors respond to distal threats (anxiety-like behaviors) compared to proximal threats (fear-like behaviors) (Mobbs et al., 2020; Perusini and Fanselow, 2015). We hypothesized that we would see the prefrontal-limbic circuit adapt as threat increases. We tested this by investigating the special case of avoidance induced by a ghost attack. During an attack, there were two possible outcomes. First, the ghost could initiate a chase, where the ghost’s speed remained constant in the direction of Pac-Man. In this case, as long as the patient turned to avoid, they had a chance to successfully escape. Second, the ghost could initiate a strike, where the ghost’s speed was increased such that even if the patient responded appropriately and fled the attack, they would be caught by the ghost and lose a life. Chase versus strike trials were counterbalanced, while, importantly, the advent of an attack was determined by the patient’s behavior in the trial. All attack trials were removed before assessing regional theta power modulations to ensure that the effects described in Figure 2 were not driven by this subset of trials.

We began by time-locking to the initiation of a ghost attack (See Figure 5b, 5f). We found similar drops in theta power in the OFC, anterior cingulate, hippocampus, and amygdala as we saw during distal-threat avoidance, despite these trials being excluded in those earlier analyses (See Figure 5f, 5e). We next turned to the HFA, where calculating the average TFR showed a striking increase in high-frequency activity after the ghost initiates an attack. However, a Bayesian mixed effects model predicting HFA in the 1 second after the attack compared to the second before the attack found that this increase was driven by only a subset of electrodes (39/127 total MFG contacts). Given that the effect within this population of electrodes was strong enough to dominate our TFRs (See Figure 5b), we wanted to understand if these electrodes were anatomically distinct. We found that the majority of the attack-activated electrodes were localized to the right hemisphere (84%; See Figure 5c). When we included hemisphere as an interaction in our HFA model before and after ghost attack, we found a significant interaction between hemisphere and attack, where right MFG contacts resulted in a significant elevation of HFA in the second after an attack began (estimate: -0.16 95% CI [-0.29 - -0.04]; See Figure 5d). To characterize what was driving HFA during the attack, we limited our analyses to the contacts in the right MFG. We hypothesized that HFA would correlate with the level of threat, where activation would decrease if the patient realized they could successfully escape. To test this hypothesis, we ran a Bayesian mixed effects model where time was used to predict HFA on the trial level. We included an interaction term for trial type (chase or strike trials), where we predicted chase trials to have a negative relationship between time and HFA as patients realized they would make a successful escape. As predicted, we found a main effect of trial type, where chase trials had decreased HFA compared to strike trials (Estimate: -0.18 95% CI [-0.21 - -0.11]) and a significant interaction with time where chase trials showed dropping HFA as patients neared the end of the trial (estimate: -0.12 95% CI [-0.17 - -0.08]; See Figure 5e). We interpret these results as evidence that high-frequency activity in the right MFG tracks the perceived threat of the ghost. Given that ghost attacks were designed to motivate the behavior on attack-free trials, we could not investigate the connectivity between the MFG during these trials as there were only an average of 58 +/-13.4 trials (across both chase and strike trials) in the patients.

**Fig. 5.**
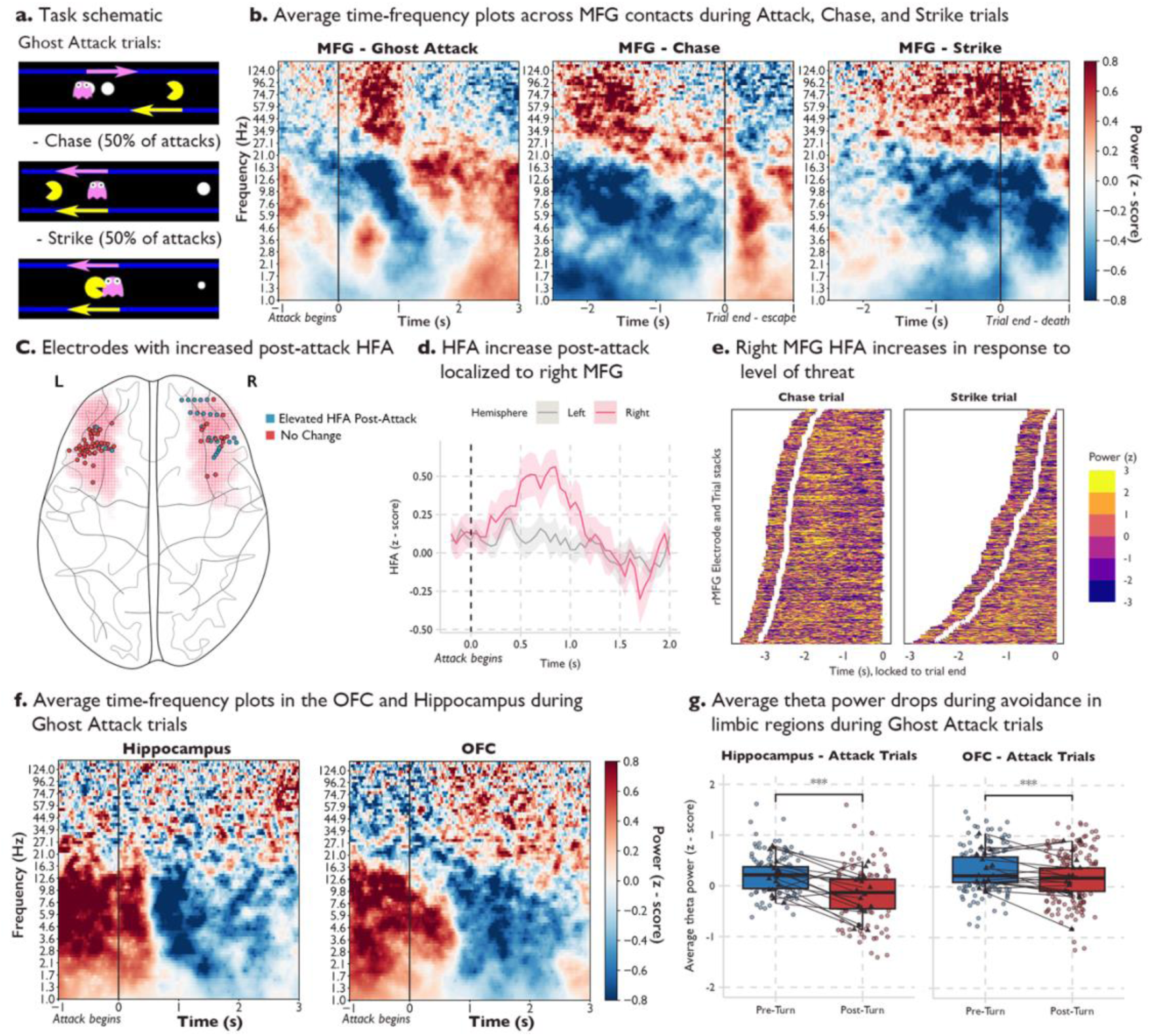
**a** Task schematic of Ghost Attack. On each trial, if the participant moved within a certain distance to the ghost, they triggered a ghost attack. Once triggered, the ghost begins moving toward the Pac-Man. On these trials, there is a 50% chance that the ghost initiates either a Chase or a Strike. On Chase trials the ghost’s speed does not change, allowing the participant the chance to respond and safely exit the trial. On Strike trials, the ghost’s speed is increased such that even if the participant appropriately responds, they will be caught by the ghost. **b** Time-frequency analyses of Ghost Attack, Chase, and Strike trials average across MFG contacts and participants. Red (blue) indicates increases (decreases) in z-scored power. The left-most plot is time-locked to the onset of a ghost attack and includes all attack trials. The middle plot is time-locked to the end of the trial and includes only Chase trials. The right-most plot is time-locked to the end of the trial and includes only Strike trials. **c** Anatomical localization of the increase in HFA after the onset of a ghost attack. Pink shading indicates the MFG, dots indicate contacts, color indicates if a given contact had significantly elevated HFA post-attack. **d** Average HFA during a ghost attack in the left MFG and right MFG contacts. Y-axis is z-scored power and the x-axis is time, locked to the onset of the attack. Color indicates hemisphere and shading indicates the standard error of the mean across participants. **e** Trial and electrode stacks of right MFG contacts during Chase trials (left plot) and Strike trials (right plot). Brighter colors indicate higher power and darker colors indicate lower power. White dots indicate the onset of the ghost attack and the x-axis is time-locked to the end of the trial. **f** Time-frequency analysis of Ghost Attack trials in the OFC and Hippocampus. Red (blue) indicates increases (decreases) in z-scored power. The X-axis is time-locked to the onset of the ghost attack. **g** Average theta power in the 1500ms before and after turning in response to onset of ghost-attack. Dots indicate average theta power in electrode across trials, triangles indicate average theta power across electrodes within a participant. Significance was assessed using Bayesian mixed effects models with random effects of participant and electrode and *** indicate the credible interval did not include 0.

## Discussion

We provide evidence for a dynamic cortico-limbic network supporting approach and avoidance. We identified approach and avoidance-dependent modulations in theta power, where theta power was elevated during the approach window in the hippocampus, amygdala, OFC, and ACC. In contrast, the middle frontal gyrus was characterized by different motifs in both the theta band and HFA. We also found increased connectivity in the prefrontal-limbic circuit, where we found theta-band coherence both across the four limbic regions, but also between the MFG, OFC, and ACC. This theta coherence increased during the approach period, as reward, threat, and conflict increased, but fell as patients turned to avoid the ghost and end the trial. We additionally found that theta oscillations in the OFC and ACC were driven by both MFG and amygdala activity, suggesting that the OFC and ACC may integrate information from both regions. The degree of synchrony between the nodes in the circuit correlated with trial-dependent changes in approach behavior, where higher theta synchrony between multiple regions, but especially with the OFC, correlated with *longer* approach times, but increased high-frequency synchrony between the hippocampus and MFG correlated with *shorter* approach times. Finally, during attack trials when the threat on a given trial shifted from distal to proximal, we found high-frequency activity in the right MFG tracked the level of threat. Together, these findings support engagement of multiple components of the prefrontal-limbic network during anxiety-inducing behavior assessed with our approach-avoidance conflict task.

Low-frequency theta oscillations are ideal for coordinating the neural activity between distal regions (Fries, 2015, 2005; Helfrich and Knight, 2016; Phillips et al., 2014). Oscillations temporally coordinate the activity between two regions to facilitate information transfer (Canolty and Knight, 2010; Yuste, 2015). In the absence of long-range synchrony, inputs may arrive at random phases of the excitability cycle in a connected region hindering effective communication between regions. Theta coherence has been proposed as a cross-region mechanism of cognitive processing and the degree of subcortical-prefrontal theta synchrony correlates with behavioral performance in a wide range of attention, learning, and working-memory tasks (Backus et al., 2016; Harris and Gordon, 2015; Johnson et al., 2017; Watrous et al., 2013). Our results support a key role for theta synchrony during approach-avoidance. We found many pairs of electrodes across the prefrontal-subcortical circuit with theta band synchrony peaking at the final decision to avoid– when the levels of threat and conflict were the highest. The prefrontal cortex is thought to have a particular role in coordinating the oscillatory activity underlying cognitive control and goal-directed behavior (Helfrich and Knight, 2016). Furthermore, in the context of approach-avoidance, Amemori et al 2024 reported that reduced connectivity in a PFC-limbic circuit results in more avoidant behavior in non-human primates (Amemori et al., 2024b). In line with these hypotheses, we found that the MFG drove theta oscillations in both the OFC and ACC, and theta synchrony across many nodes in the prefrontal-limbic theta network correlated with longer approach times in the task, meaning that patients took on more risk to receive increased reward in those trials. Importantly, while Amemori focused on alpha oscillations in a restricted set of limbic regions, they defined the alpha band as between 5-13Hz, and reported peaks at 7Hz, which overlaps with our 3-8Hz theta band definition for this study.

We also found that HFA synchrony had differential effects on behavior depending on regional pairing. While the OFC-MFG and OFC-ACC HFA synchrony correlated with longer approach times, hippocampus-MFG HFA synchrony predicted shorter approach times. This indicates that patients were more likely to forgo reward in favor of lower risk when MFG and hippocampus had similar HFA profiles. Recent work suggests that high-frequency power correlations may be reflective of functional, but not direct, connectivity (Helfrich and Knight, 2016; Hipp et al., 2012), meaning that the similar HFA profiles in these regions may result from the coordination by a third region, rather than the result of direct neuronal projections between the MFG and hippocampus. Supporting this interpretation, we found lower theta synchronization between the hippocampus and MFG. Instead, we observed that the OFC and ACC share high theta coherence with both the subcortical regions and the MFG, suggesting that these regions may be integrating the information across distinct neural circuits. This has been hypothesized for the ACC under reward and threat processes (Clarke et al., 2015; Wallis et al., 2019), where ACC-Amygdala connectivity is proposed to track uncertain threat (Kirk et al., 2022), while PFC-ACC connectivity may relate to emotion regulation (Etkin et al., 2015; Rolle et al., 2022). Our directionality analyses support these reports with both amygdala and lateral prefrontal activity driving theta oscillations in the OFC and ACC.

The hippocampus is an important node within this circuit and is a key area of study in approach-avoidance paradigms in rodents (Bryant and Barker, 2020; Gray, 1982; for recent reviews, see Ito and Lee, 2016; Kimura, 1958). Recent studies in both rodents and humans have proposed that threat information is propagated from the hippocampus to the prefrontal cortex via theta during approach-avoidance (Abivardi et al., 2020; Padilla-Coreano et al., 2019). Padilla-Coreano et al 2019 found that disrupting hippocampus to mPFC theta synchrony reduced avoidance but increasing synchrony increased avoidance behavior in mice. Further analyses revealed that it was not the synchrony itself that determined the avoidance behavior, but the degree of information transmission from the hippocampus to the mPFC that biased the animals toward avoidance. In our analyses, we found that theta synchrony correlated with *longer* approach times but hippocampus-MFG high-frequency synchrony resulted in earlier avoidance. Thus, while similar representations of threat information in the hippocampus and MFG may lead to early avoidance, other types of information, such as reward or uncertainty, may also propagate through the circuit. Accordingly, other region pairs, such as the OFC-ACC and OFC-MFG had positive relationship between HFA synchrony and approach times.

Finally, fitting with a model of approach-avoidance behavior where distal and uncertain threat results in anxiety-like behavior while proximal and certain threat induces fear behaviors (Mobbs et al., 2020; Perusini and Fanselow, 2015), we see rapid changes in the cortical-subcortical circuit during ghost attack. Specifically, the time-frequency plots were dominated by a rapid increase in high-frequency activity in the right MFG during ghost attack. Previous work has reported evidence of left/right asymmetry in the lateral PFC, which includes the MFG, especially in the context of emotional regulation (Brunoni et al., 2017; White et al., 2023). It has been shown that the right dlPFC is activated during periods of unpredictable threat, and this increased dlPFC activation is associated with reduced anxiety (Drevets and Raichle, 1992). These studies suggest that the right dlPFC enacts emotional regulation by providing top-down control on the limbic regions. In our study, the high-frequency activity during the acute threat period may provide a control signal as patients try to regulate their escape in the face of a near-term threat. Once patients realize they will make a successful escape the need for this control drops and is accompanied by a rapid decrease in high-frequency activity.

While individual regional contributions to approach-avoidance behavior have been identified across human, NHP, and rodent studies, this is the first study in humans to characterize the theta-mediated circuit dynamics that enable these regions to function in concert in real-time. Human fMRI and NHP research have highlighted the crucial role of widespread prefrontal involvement, while rodent studies have implicated mPFC-subcortical theta synchrony in regulating approach and avoidance behaviors. Our findings integrate these literatures by demonstrating that theta coherence links both subcortical structures with prefrontal regions, but also links both the cognitive and limbic regions within the prefrontal cortex. The circuit architecture identified here suggests that approach-avoidance behavior is governed by hierarchically organized theta-based interregional networks. For instance, the hippocampus and amygdala exhibit robust theta coherence, but amygdala signals propagate to both the OFC and ACC, allowing integration with input from the MFG. Importantly, by examining this behavior in real time, we show that network strength increases as the individual reaches the decision point between approach and avoidance, suggesting that this circuit is most engaged when threat and conflict intensifies.

There are caveats to this research. First, our study was conducted on patients with epilepsy undergoing clinical evaluation, raising the question of the generalizability of our findings. To address this, we undertook extensive efforts to only test patients fully alert and cooperative and excluded electrodes near seizure foci or contaminated by artifacts. We also show similar behavioral patterns across our patients and an online sample of 191 participants (See Figure 1, panels c-f). Also, it is important to note that electrodes were placed based on the clinical needs of the patient. As such, coverage varied across patients and some effects may have been missed due to reduced coverage. We attempted to control for this by using hierarchical models that accounted for within-patient and within-region correlations.

Similarly, coverage limitations prevented us from analyzing subregion effects, as between the anterior and posterior hippocampus. The rodent literature indicates that the ventral hippocampus corresponding to the anterior hippocampus in humans is involved in approach-avoidance as we observed (Bryant and Barker, 2020; Ito and Lee, 2016). However, we were not powered to test for anterior versus posterior hippocampal effects.

In conclusion, we used intracranial recordings and a continuous-choice approach-avoidance task in humans to characterize prefrontal-limbic neural activity during anxiety-inducing decision-making. We found broad regional involvement with a particular focus on theta power modulations and coherence supporting a distributed prefrontal-limbic network underlying approach-avoidance. The findings contribute to a circuit understanding of psychiatric disorders, especially those characterized by maladaptive avoidance, such as generalized anxiety disorder, agoraphobia, and social anxiety disorder.

## Methods

### Patients

We recorded intracranial signals from twenty (10) female adult patients (mean age 27.25 +/- 12.04) who underwent intraoperative neurosurgical treatment for pharmacoresistant epilepsy. All patients were implanted with depth electrodes to localize the seizure onset zone for eventual surgical resection. We selected patients with implantations in any of the following regions: hippocampus, amygdala, OFC, anterior cingulate, and middle frontal gyrus. The electrode placement was dictated solely by the patient’s clinical team. See Supplemental Figure 1 for patient specific electrode implantation. The recordings took place at the following hospitals: Loma Linda University Medical Center (n = 6), Barnes-Jewish Hospital in St. Louis (n = 12), and the St. Louis Children’s Hospital (n=2). We additionally collected 191 participants from the online recruitment platform, Prolific. The online participants were paid a set amount for their participation in the research, along with a bonus that correlated with task performance.

Participants were given thorough instructions along with a short quiz to ensure comprehension of the task. All patients provided written informed consent as part of the research protocol approved by each hospital’s Institutional Review Board and by the University of California, Berkeley. Patients were tested when they were fully alert and willing to participate.

### Behavioral Task

We investigated approach-avoidance dynamics using a novel, continuous-choice approach-avoidance task based on the arcade game Pac-Man. Figure 1a demonstrates the experimental paradigm. At the beginning of each trial, Pac-Man is placed along a single corridor that runs horizontally in the center of the screen. The player makes a button press (either the left or right arrow key) to begin moving along the corridor. They can either choose to move towards the *end* of the corridor where they can exit to the next trial or they can move towards the *center* of the corridor whereby the patient encounters both potential gains (eating “dots”, resulting in points) and potential losses (ghost attack, resulting in loss of a life). Patients could choose to eat up to 5 dots on each trial, but they could also choose to exit the trial at any time. The dots vary in size indicates their worth (large dots = 20 points, small dots = 10 points). However, importantly, both being chased and being caught by the ghost are determined by the player’s distance to the ghost. The closer the participant chooses to go to the ghost, the higher the likelihood that the ghost will initiate an attack. Thus, choosing to acquire more points always increased the risk of being eaten by the ghost. The probability of an attack is set by a beta distribution on the possible distances from Pac-Man to the ghost. If the ghost initiates an attack, there is a 50% chance the ghost will initiate a Chase and a 50% chance the ghost will initiate a strike. In a chase trial, the ghost begins to move towards the Pac-Man but the speed is unchanged, allowing the participant to make a button response to turn and exit the trial safely. In strike trials, the ghost’s speed is modulated such that even if the participant responds appropriately, they will be caught by the ghost and lose a life. 20% of trials were conflict-free, where the player was free to collect dots without the risk of losing a life, though they did have to return to exit the corridor on the side closest to where the trial began to ensure similar movement across conflict and conflitc-free trials. The game is played as a series of mini-games, which end when either 1) Pac-Man loses all three lives or 2) a block of twenty trials are completed. At the end of each minigame, the score was reset and the patient was prompted to begin a new game. Direction, reward, location of the ghost, location of Pac-Man, and the ghost’s starting direction are counterbalanced across trials. Patients were shown and read instructions prior to playing a short practice and were explicitly told that 1) the closer they got to the ghost the higher the chance of attack and 2) that to achieve the highest possible score, they should balance the risks of being caught by the ghost and the rewards of getting as many dots as possible. The game itself was composed of 240 trials. A full experimental run typically lasted approximately 20-25 minutes. Stimulus presentation was operated by JavaScript running in a Chrome browser.

### Data acquisition

Electrophysiological data were recorded using BCI2000, an open-source software, at each clinical site (Schalk et al., 2004). This system synchronized the Pac-Man task with LFPs, eye tracking, and behavior (Pac-Man movement) in a single data stream which supports easy data pooling across sites and data sharing with the community (Milsap et al., 2019). The sampling rate at Barnes-Jewish Hospital was 2000Hz, while at Loma Linda the sampling rate was 512Hz.

#### Anatomical reconstructions

We used an anatomical data processing pipeline to localize electrodes from a pre-implantation MRI and a post-implantation CT scan (Stolk et al., 2018). The MRI and CT images were aligned to a common coordinate system and fused with each other using a rigid body transformation. We then compensated for brain shifts caused by the implantation surgery. A hull of the patient brain was generated using the Freesurfer analysis suite (Dale et al., 1999). Electrodes were then classified by a neurologist according to the anatomical location within each patient’s anatomical space. Only electrodes confirmed to be in the regions of interest were included in the analysis. For illustration purposes only, we converted patient-space electrodes into Montreal Neurological Institute (MNI) coordinates using volume-based normalization. Figure 1f shows all the electrodes used in the analysis.

### Data preprocessing

Offline, continuous data were low-pass filtered at 150Hz, and notched filtered at 60Hz and its harmonics up to 120Hz to remove line noise. Electrodes were re-referenced using a bipolar reference scheme. We visually identified and removed channels with excessive noise or with poor contact and each dataset was visually inspected by a neurologist to remove electrodes exhibiting epileptic spiking activity and epochs where spiking spread from the primary epileptic site. Data were epoched based on the time point of interest, either trial onset, turnaround choice, or trial end. Oscillatory and high-frequency activity was extracted at these timepoints using Morlet wavelets where the power of 80 log-spaced frequencies ranging from 1-150 was calculated using an increasing number of cycles, specifically 80 log-spaced cycles ranging from 2-30. Data preprocessing was carried out in Python using the MNE package (Larson et al., 2024).

### Power estimates

Theta-band and high-frequency power estimates were extracted from the TFR for all statistical analyses. Power was log-transformed and each specific frequency was z-scored using an ITI baseline. Each frequency between 3-8Hz was then averaged, resulting in a theta-band power estimate. For the high-frequency activity estimate, we followed a similar approach but averaged the individual frequencies in 20Hz overlapping subbands (e.g., 70-90Hz, 80-100Hz, … 130-150Hz) before averaging the power in each subband for a final estimate of high-frequency activity.

For the cross-correlation analysis, we needed both power and phase information within the theta-band. We therefore recalculated theta by bandpass filtering between 3-8Hz using a zero-phase FIR filter with a Hamming window on all electrode pairs with elevated theta coherence. Data were z-scored to the ITI, time-locked to movement-onset, detrended, and only data during approach were included in this analysis.

### Connectivity Measures - Coherence & Synchrony

Data epoched to the turnaround choice were extracted between the 2500 milliseconds before and after the turnaround using Python-MNE (Larson et al., 2024). Coherence measures were computed over time but across trials for every combination of electrodes across the regions of interest using the package MNE-connectivity. Data were resampled to 100Hz to speed computation time. Spectrum estimation was done using Morlet Wavelets using the same parameters as above, but only within the theta-band (3-8Hz). Three different coherence measures were calculated to ensure our results were not biased by the choice of metric: imaginary coherence, pairwise phase consistency, and the debiased estimator of the squared, weighted Phase Lag Index. To identify electrodes with a significantly elevated level of theta coherence, the true coherence metrics were compared, timepoint-by-timepoint, to a shuffled null distribution where the trials in one region were shuffled 1000 times. P values were calculated across timepoints and FDR-corrected to account for multiple comparisons. Electrode pairs were included in subsequent analyses if there were 100 milliseconds with significantly elevated coherence. Note, for imaginary coherence we calculated the absolute value of the imaginary coherence before calculating the P values.

As trial-level estimates of coherence were unstable, when testing for associations with trial-level behavior we used the correlation between two electrodes’ power envelopes. For this analysis, we time-locked theta power to the final decision to avoid and filtered out the beginning of the trial when Pac-Man was stationary. We then calculated the Pearson correlation trial-by-trial for every combination of electrodes across the regions of interest. We applied this approach for both the theta-band power and for high-frequency activity estimates.

### Spectral Granger Analysis

Spectral Granger causality (GC) quantifies the directionality of connectivity within specific frequency bands. Specifically, we used a State-Space formulation of Granger Causality as implemented in Python-MNE (Larson et al., 2024), which can be thought of as a more versatile and robust formulation to previous parametric approaches (Barnett and Seth, 2015). To improve computational efficiency, data were resampled to 100 Hz. Frequency estimation between 3–8 Hz was performed using Morlet wavelets with the same parameters as above. Spectral Granger causality scores were computed using 12 lags, which in spectral State-Space GC acts as a spectral smoothing parameter across the frequencies. Granger values were then averaged across the 3–8 Hz frequency range. Following the approach of Haufe and colleagues (Haufe et al., 2013) we additionally contrasted the scores with net GC scores calculated on the reversed-times series, as flipping the direction of the signal should similarly flip the estimate of directed connectivity. Thus, our final net spectral Granger scores were calculated as:

Net Granger scores were calculated for each timepoint in the final 2500ms before the decision to avoid in all electrode pairs with significant levels of theta coherence. Only timepoints in the last 1500ms of approach were analyzed to avoid edge artifacts. These timepoints were compared to a permuted null distribution where trial labels from one region were shuffled 200 times. The absolute value of the Net Granger scores were used to calculate permuted p values compared to this distribution. We then FDR-corrected for the number of timepoints and included electrode pairs in our subsequent analysis with 500ms of significantly elevated net Granger scores. The net Granger scores from these electrode pairs were the fed into our directionality tests (see below).

### Cross-correlation Analysis

To further validate our directionality findings, we additionally conducted a cross-correlation analysis within the theta-band. After calculating bandpass filtering in the the theta-band (see Power Estimation for details), we then concatenated all approach windows across the trials into one sequence per pair of electrodes. We next calculated a cross-correlation function with 25 lags corresponding to 250ms on the concatenated theta signal. We then created a permuted distribution by shuffling the trial labels 200 times, while keeping the trial lengths constant to ensure that the discontinuities in the true concatenated signal were equivalent in each of the permuted samples. To assess if there was a significant correlation between two electrodes, we compared the correlation value from the true pair to the permuted distribution. If the correlation was in the top 5% of the permuted distribution, its best lag was included in our directionality test (see below).

### Bayesian Linear Mixed Effects Models

Effects were assessed using Bayesian mixed effects models that always included hierarchically grouped random effects of patient and electrodes. The benefit of these models is they provide a single test of our hypotheses, eliminating the need to correct for the number of electrodes or develop second-level tests of how results varied across patients, while appropriately accounting for the correlations within electrodes and patients. This hierarchical random effects structure meant fitting models at times with over 3000 random effect levels. Bayesian models fit with MCMC sampling allowed us to appropriately and effectively fit this complex model structure.

Across all the Bayesian mixed effects models, we used a standard set of weakly informative priors. For intercepts, the prior was set to the normal distribution, centered on 0, with a standard deviation of 5. For beta coefficients, the prior was set to the normal distribution, centered on 0, with a standard deviation of 2. For the standard deviation of the random effects, we used an exponential distribution with a positive arrival rate of 1. Finally, we set a prior on the correlation between random effects using the Lewandowski-Kurowicka-Joe distribution with the shape parameter of 2. All models used 4 chains and were initially fit using 5000 iterations of which 1000 were used as warmup. If there were indications that the models did not converge, we increased the number of iterations to a max of 10000 iterations, of which 5000 were warmup. Rhat metrics, effective sample size and trace plots were used to confirm model convergence and high resolution; all Rhat values for estimates were equal to or below 1.01 and all effects of interest had Rhat values equal to 1.0. Effective samples were all greater than 1000. Posterior predictive checks on the mean and standard deviation were completed across all models (See Supplemental Table 3, 4 & 7). All models were fit using the package “brms” in R (Bürkner, 2017).

### Directionality Analyses

To assess whether net Granger causality values or cross-correlation time exhibited a consistent directional trend across patients, we employed a hierarchical Bayesian model using the brms package in R. The model estimated the central tendency of net Granger values while accounting for patient- and electrode-level variability. We used weakly informative priors (see above for details). To quantify the probability that net Granger values or time lags were consistently positive across patients, we computed the Probability Positive (P+), defined as the proportion of posterior samples where the estimated effect exceeded zero. This metric provides an intuitive, probabilistic measure of directional certainty, with P+ values closer to 1 indicating strong evidence for consistent directionality.

### Data and code availability

All data reported in this paper will be shared by the lead contact upon request. Original code will be deposited on Zenodo for public download. Any additional information required to reanalyze the data reported in this paper is available from the lead contact upon request.

## Author contributions

B.R.S., R.T.K, and M.H. designed the experiment; R.T.K., O.K.M., J.T.W., P.B, M.D., and J.J.L. supervised data collection; O.K.M., J.T.W., P.B, M.D., J.J.L and B.R.S. collected the data; B.R.S. analyzed the data; J.O. analyzed online behavioral task data; B.B and T.M. helped to preprocess the data and edited the manuscript. B.R.S, M.H., and R.T.K. interpreted the data; B.R.S. wrote the manuscript; and M.H. and R.T.K. supervised the project.

Funding: NSF GRFP (BRS); NIH/NINDS R01 NS-021135 (RTK); NIH/NIBIB P41 EB-018783 (PB); NIH/NINDS R01 EB-026439 (PB); NIH/NINDS U24 NS-109103 (PB); NIH/NINDS U01 NS-108916 (PB); NIH/NIMH R01 MH-120194 (JTW); McDonnell Center for Systems Neuroscience (PB); Fondazione Neurone (PB), American Epilepsy Society (JTW)

