## SupplementalMaterials for "Circuit dynamics of approach-avoidance conflict in humans"

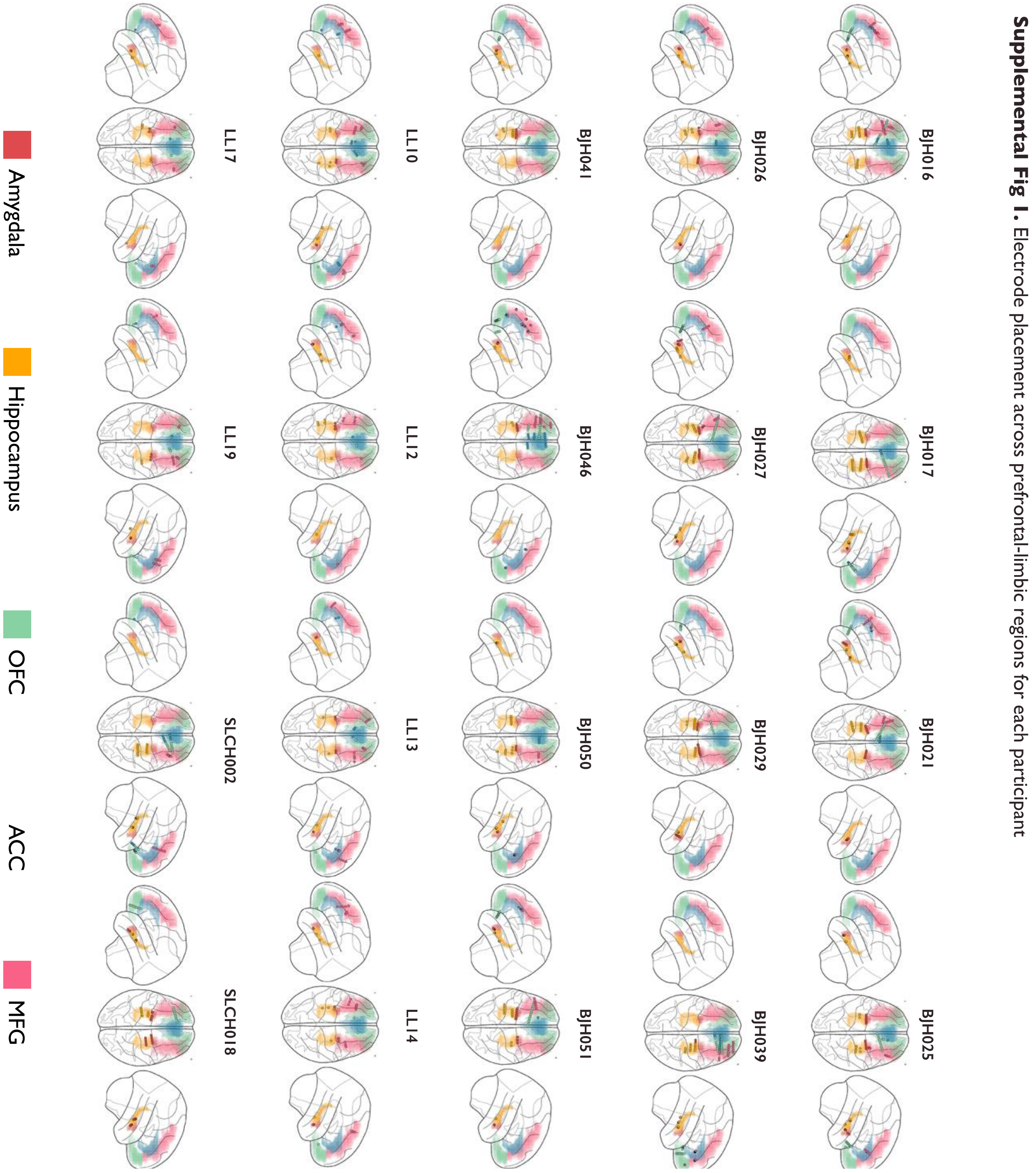

***Supp Fig. 1.*** Electrode placement for each of the twenty patients. Electrodes were placed across the prefrontal and limbic regions. Colored shading indicates region and dots indicate electrode.

**Supplemental Fig 2.** Time-frequency representations of conflict-free trials, time-locked to avoidance choice in all regions

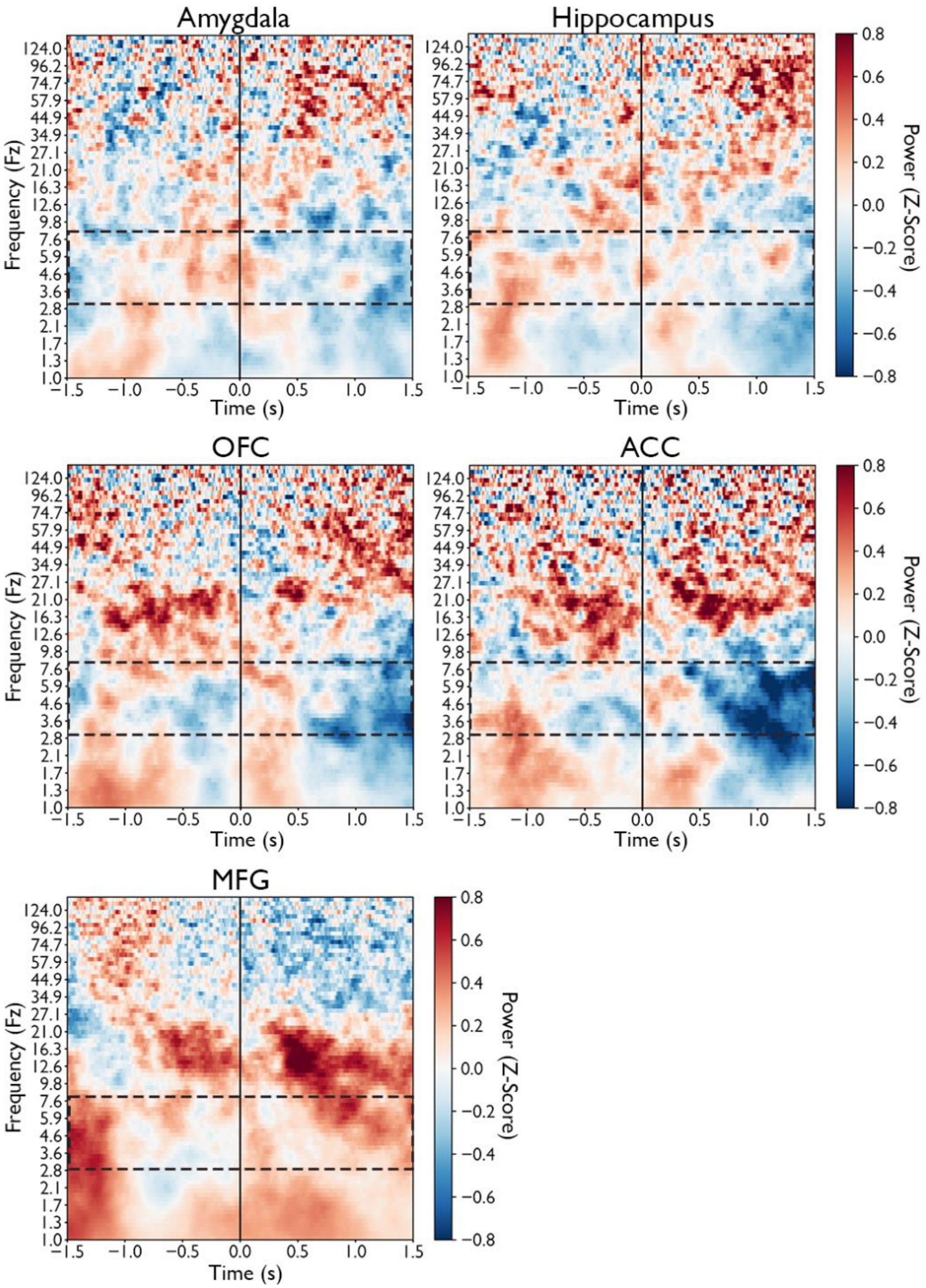

***Supp Fig. 2***. Time-frequency analysis of neural activity time-locked to the final choice to stop approaching and begin avoiding in conflict-free (reward-only, no ghost) trials across all electrodes in the limbic regions (hippocampus, amygdala, OFC and ACC) and the MFG. Red (blue) indicates increases (decreases) in z-scored power. Dotted box indicates the theta band (3-8Hz).

**Supp. Fig. 3:** Theta Oscillations in the Middle Frontal Gyrus

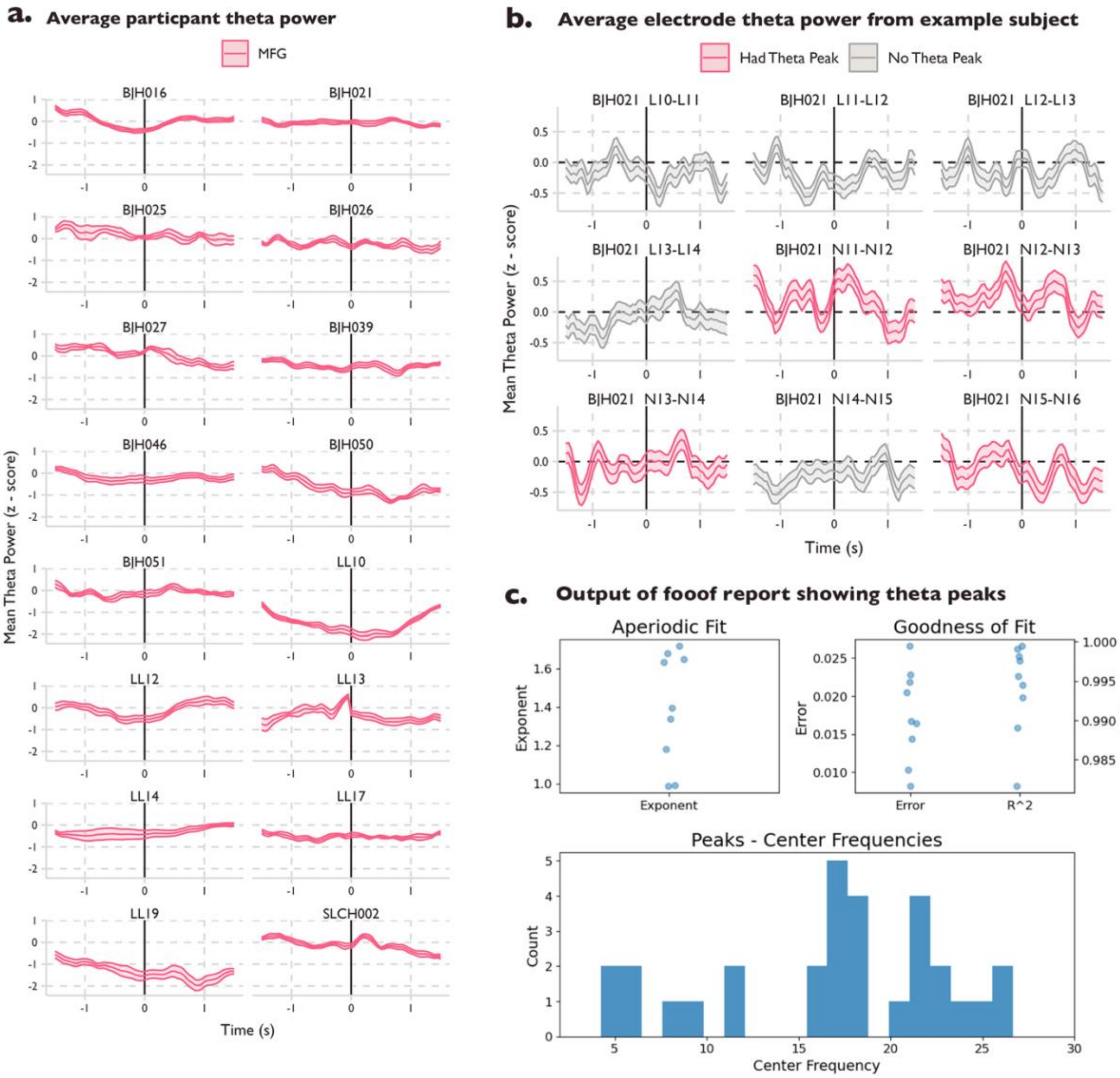

***Supp Fig. 3.*** **a** Time-course of theta power in the MFG for each patient across approach and avoidance windows. 0 indicates the time the patient stopped approaching and began avoiding. Shading is the standard error of the mean power across subjects. **b** Average time-course of theta power by electrode for an example subject. While this example subject had an average near 0 of z-scored theta power, when looking at the electrode level there are dynamics in the time-course. 0 indicates the time the patient stopped approaching and began avoiding. Shading is the standard error of the mean and the color indicates if that electrode had a peak in the theta band, as assessed by the FOOOF (Fitting Oscillations and One-Over-F) algorithm—which decomposes the power spectrum into its aperiodic (1/f) and periodic components. **c** Output for a group-level FOOOF report run on all the electrodes for the example participant in panel b. Output shows high R-squared values and low error. While peaks in the beta-band were more common, 4/12 electrodes also showed peaks in the theta-band.

**Supp Table 1.** Percentage of electrode pairs with significantly elevated theta coherence for each region.

| Mean Percentage of Sig. Elec Pairs | | | | |
|---|---|---|---|---|
| | Mean | SD | Min | Max |
| *Hippocampus* | | | | |
| Imaginary Coherence | 36% | 16% | 10% | 68% |
| Pairwise Phase Consistency | 46% | 22% | 15% | 88% |
| Phase Lag Index | 37% | 17% | 6% | 67% |
| *Amygdala* | | | | |
| Imaginary Coherence | 36% | 12% | 17% | 62% |
| Pairwise Phase Consistency | 52% | 19% | 17% | 83% |
| Phase Lag Index | 38% | 14% | 17% | 67% |
| *OFC* | | | | |
| Imaginary Coherence | 38% | 17% | 8% | 85% |
| Pairwise Phase Consistency | 50% | 18% | 17% | 94% |
| Phase Lag Index | 40% | 17% | 13% | 88% |
| *ACC* | | | | |
| Imaginary Coherence | 45% | 16% | 20% | 75% |
| Pairwise Phase Consistency | 59% | 17% | 21% | 83% |
| Phase Lag Index | 46% | 18% | 11% | 83% |
| *MFG* | | | | |
| Imaginary Coherence | 33% | 18% | 14% | 74% |
| Pairwise Phase Consistency | 44% | 21% | 14% | 90% |
| Phase Lag Index | 34% | 18% | 14% | 76% |

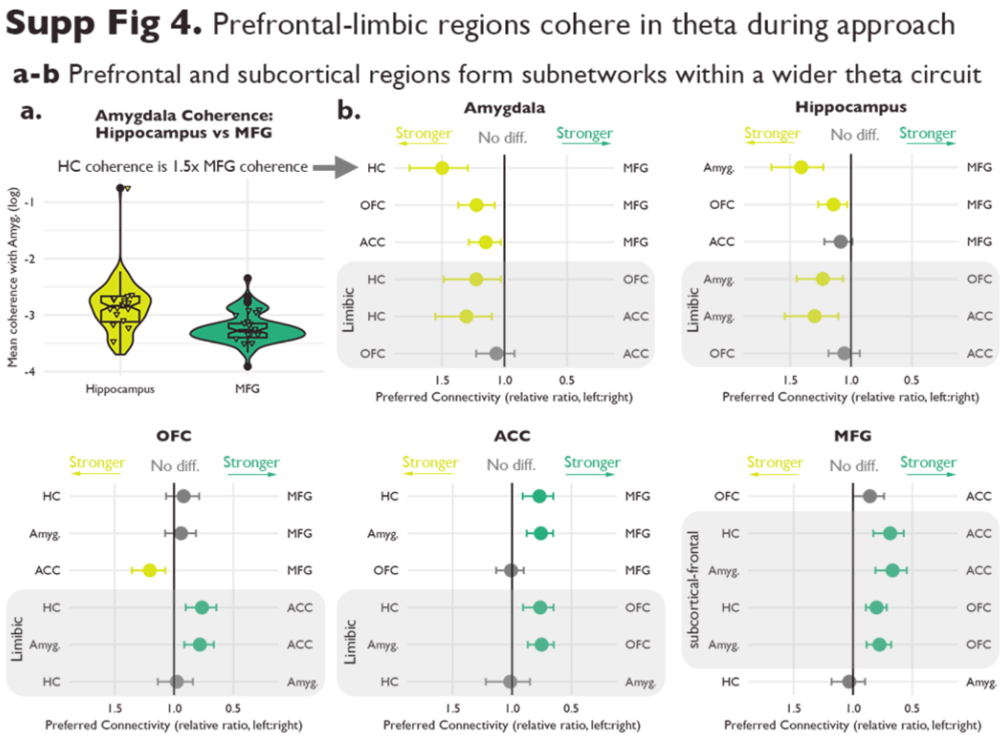

*Supp Fig. 4.* Full interregional comparison of theta-band imaginary coherence strength. Theta coherence in each region was modeled with all other regions as predictors, similar to a frequentist ANOVA. **a** Comparison of estimated theta coherence strength between the Hippocampus and MFG, within the Amygdala. X-axis is the region, y-axis if the log-scaled estimated coherence with the amygdala. This plot is summarized in the first line of the first subplot in b. **b** Each row on the y-axis compares the coherence strength between the left and right regions. Dots indicate the estimated relative ratio of preferred connectivity. Lines indicate the 95% credible interval. Grey indicates the credible interval includes 1, meaning there is no difference in coherence strength between the two regions. For the limbic subplots, grey shading highlights within limbic comparisons, while for the MFG, the grey shading highlights subcortical-frontal comparisons.

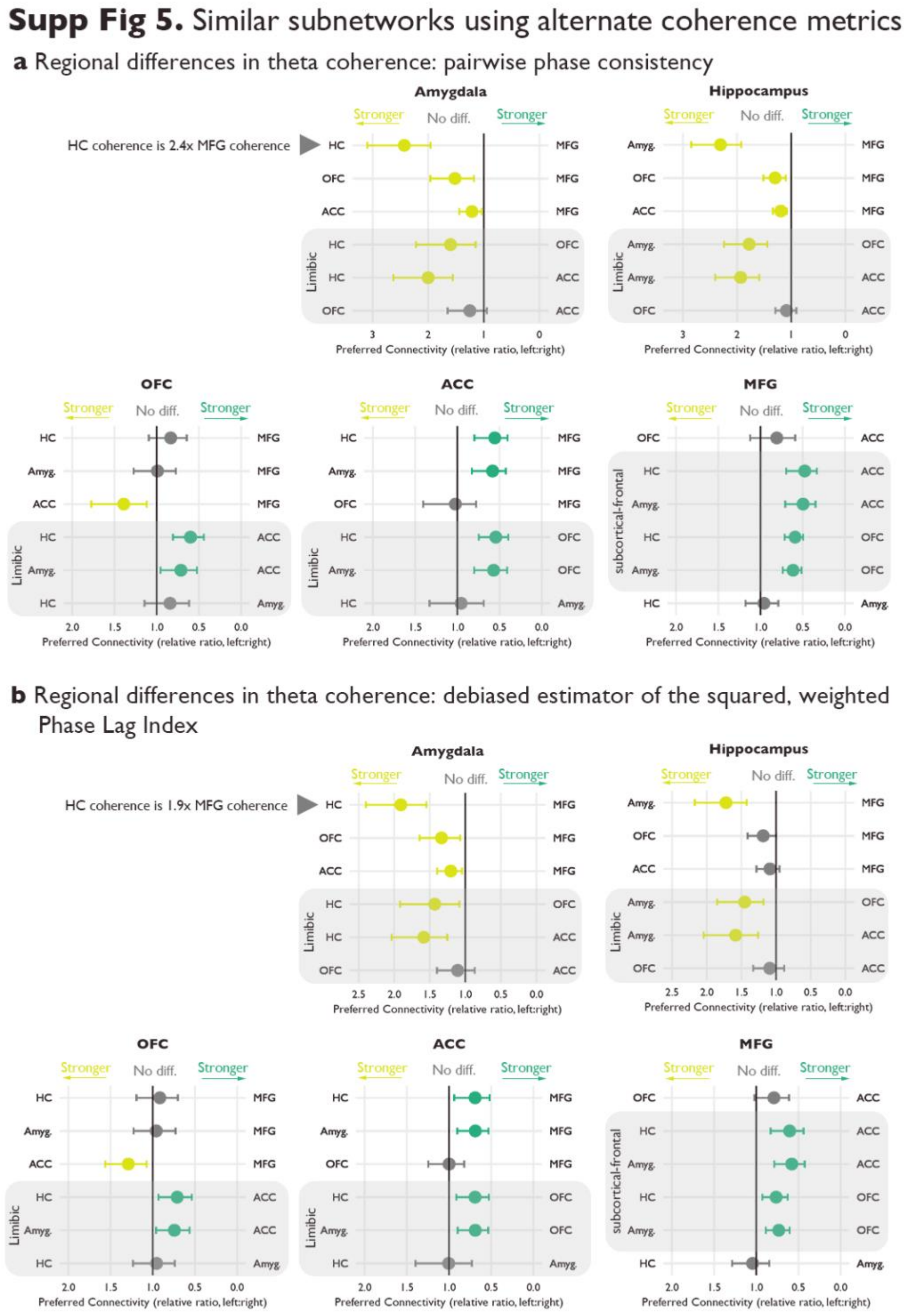

***Supp Fig. 5.*** Theta connectivity profiles using **a** pairwise phase consistency **b** phase lag index. Theta coherence in each region was modeled with all other regions as predictors. Each row on the y-axis compares the coherence strength between the left and right regions. Dots indicate the estimated relative ratio of preferred connectivity. Lines indicate the 95% credible interval. Grey indicates the credible interval includes 1, meaning there is no difference in coherence strength between the two regions. For the limbic subplots, grey shading highlights within limbic comparisons, while for the MFG, the grey shading highlights subcortical-frontal comparisons.

### **Supp. Table 2.** Full regional theta coherence results across all regions and metrics

##### Amygdala Coherence

| Metric | Mean | l-95% CI | u-95% CI | Est.Error | Rhat | Bulk_ESS | Tail_ESS |
|---|---|---|---|---|---|---|---|
| **Hippocampus - Ant.Cingulate** | | | | | | | |
| *PPC* | 2.00 | 1.56 | 2.63 | 0.13 | 1 | 6210.77 | 6843.46 |
| *PLI* | 1.58 | 1.25 | 2.04 | 0.12 | 1 | 7952.22 | 6605.36 |
| *IMCOH* | 1.30 | 1.10 | 1.55 | 0.09 | 1 | 8677.27 | 7881.79 |
| **Hippocampus - OFC** | | | | | | | |
| *PPC* | 1.60 | 1.15 | 2.22 | 0.17 | 1 | 5112.92 | 6659.38 |
| *PLI* | 1.43 | 1.08 | 1.92 | 0.14 | 1 | 6466.17 | 6274.19 |
| *IMCOH* | 1.23 | 1.03 | 1.49 | 0.09 | 1 | 7495.55 | 6934.92 |
| **Hippocampus - MFG** | | | | | | | |
| *PPC* | 2.43 | 1.96 | 3.10 | 0.12 | 1 | 5590.09 | 5962.05 |
| *PLI* | 1.91 | 1.55 | 2.40 | 0.11 | 1 | 6942.64 | 5960.14 |
| *IMCOH* | 1.50 | 1.29 | 1.76 | 0.08 | 1 | 7582.33 | 6987.06 |
| **Ant.Cingulate - OFC** | | | | | | | |
| *PPC* | 0.80 | 0.61 | 1.05 | 0.14 | 1 | 6037.59 | 7399.08 |
| *PLI* | 0.90 | 0.72 | 1.15 | 0.12 | 1 | 7236.54 | 8111.59 |
| *IMCOH* | 0.94 | 0.82 | 1.09 | 0.07 | 1 | 8807.66 | 9204.87 |
| **Ant.Cingulate - MFG** | | | | | | | |
| *PPC* | 1.21 | 1.04 | 1.44 | 0.08 | 1 | 8890.59 | 7189.73 |
| *PLI* | 1.20 | 1.04 | 1.40 | 0.07 | 1 | 13269.23 | 8905.61 |
| *IMCOH* | 1.15 | 1.03 | 1.28 | 0.06 | 1 | 11399.59 | 8496.83 |
| **OFC - MFG** | | | | | | | |
| *PPC* | 1.52 | 1.18 | 1.96 | 0.13 | 1 | 4942.21 | 7048.68 |
| *PLI* | 1.33 | 1.07 | 1.64 | 0.11 | 1 | 6436.95 | 7849.50 |
| *IMCOH* | 1.22 | 1.08 | 1.37 | 0.06 | 1 | 6426.08 | 7442.93 |

**Random effect of subject (20 levels)**

| Metric | Estimate | Est.Error | l-95% CI | u-95% CI | Rhat | Bulk_ESS | Tail_ESS |
|---|---|---|---|---|---|---|---|
| sd(Intercept) | | | | | | | |
| PPC | 0.34 | 0.09 | 0.19 | 0.53 | 1 | 4568.84 | 6461.53 |
| PLI | 0.27 | 0.08 | 0.13 | 0.45 | 1 | 4484.15 | 4757.39 |
| IMCOH | 0.15 | 0.05 | 0.05 | 0.26 | 1 | 2836.47 | 2572.59 |
| sd(OFC) | | | | | | | |
| PPC | 0.42 | 0.12 | 0.20 | 0.69 | 1 | 4383.08 | 4200.46 |
| PLI | 0.34 | 0.10 | 0.18 | 0.56 | 1 | 5745.05 | 6528.40 |
| IMCOH | 0.15 | 0.07 | 0.02 | 0.29 | 1 | 2527.43 | 2199.27 |
| sd(Ant.Cingulate) | | | | | | | |
| PPC | 0.11 | 0.09 | 0.00 | 0.33 | 1 | 5346.63 | 5663.81 |
| PLI | 0.11 | 0.08 | 0.00 | 0.31 | 1 | 5448.11 | 5882.55 |
| IMCOH | 0.08 | 0.06 | 0.00 | 0.24 | 1 | 4053.52 | 4867.05 |
| sd(Hippocampus) | | | | | | | |
| PPC | 0.23 | 0.15 | 0.01 | 0.57 | 1 | 2292.98 | 4582.82 |
| PLI | 0.27 | 0.17 | 0.01 | 0.66 | 1 | 1730.82 | 4137.30 |
| IMCOH | 0.20 | 0.12 | 0.01 | 0.46 | 1 | 1664.83 | 2716.07 |
| cor(Intercept,OFC) | | | | | | | |
| PPC | -0.16 | 0.27 | -0.65 | 0.40 | 1 | 5376.21 | 7510.14 |
| PLI | -0.05 | 0.28 | -0.56 | 0.51 | 1 | 6194.74 | 7483.67 |
| IMCOH | -0.12 | 0.33 | -0.69 | 0.56 | 1 | 5806.37 | 7809.13 |
| cor(Intercept,Ant.Cingulate) | | | | | | | |
| PPC | 0.11 | 0.36 | -0.62 | 0.75 | 1 | 14459.04 | 8634.25 |
| PLI | 0.16 | 0.36 | -0.59 | 0.78 | 1 | 13826.58 | 8132.39 |
| IMCOH | -0.01 | 0.37 | -0.69 | 0.69 | 1 | 13902.68 | 8542.65 |
| cor(OFC,Ant.Cingulate) | | | | | | | |
| PPC | 0.09 | 0.37 | -0.64 | 0.74 | 1 | 13434.11 | 9308.95 |
| PLI | -0.03 | 0.36 | -0.71 | 0.65 | 1 | 14968.28 | 8133.88 |
| IMCOH | 0.05 | 0.37 | -0.66 | 0.73 | 1 | 12709.59 | 9030.68 |
| cor(Intercept,Hippocampus) | | | | | | | |
| PPC | 0.19 | 0.34 | -0.51 | 0.78 | 1 | 9215.03 | 8626.87 |
| PLI | 0.05 | 0.34 | -0.61 | 0.68 | 1 | 9658.15 | 8272.69 |
| IMCOH | 0.00 | 0.33 | -0.63 | 0.63 | 1 | 7818.30 | 8690.99 |
| cor(OFC,Hippocampus) | | | | | | | |
| PPC | -0.10 | 0.34 | -0.72 | 0.57 | 1 | 8469.50 | 8508.68 |
| PLI | -0.06 | 0.33 | -0.69 | 0.59 | 1 | 8507.00 | 8091.84 |
| IMCOH | -0.23 | 0.34 | -0.80 | 0.50 | 1 | 4624.24 | 6789.36 |
| cor(Ant.Cingulate,Hippocampus) | | | | | | | |
| PPC | 0.08 | 0.38 | -0.65 | 0.76 | 1 | 6570.21 | 8065.12 |
| PLI | 0.13 | 0.37 | -0.63 | 0.77 | 1 | 4864.30 | 7592.56 |
| IMCOH | 0.06 | 0.37 | -0.66 | 0.73 | 1 | 5285.86 | 8610.10 |

##### Hippocampus Coherence

| Metric | Mean | l-95% CI | u-95% CI | Est.Error | Rhat | Bulk_ESS | Tail_ESS |
|---|---|---|---|---|---|---|---|
| **Ant.Cingulate - OFC** | | | | | | | |
| *PPC* | 0.92 | 0.77 | 1.10 | 0.09 | 1 | 4154.52 | 6070.33 |
| *PLI* | 0.92 | 0.75 | 1.13 | 0.10 | 1 | 5575.60 | 6372.68 |
| *IMCOH* | 0.95 | 0.84 | 1.08 | 0.06 | 1 | 7121.59 | 8282.35 |
| **Ant.Cingulate - Amygdala** | | | | | | | |
| *PPC* | 0.52 | 0.42 | 0.63 | 0.11 | 1 | 5411.76 | 6959.40 |
| *PLI* | 0.63 | 0.49 | 0.79 | 0.12 | 1 | 5306.36 | 6751.61 |
| *IMCOH* | 0.77 | 0.65 | 0.91 | 0.08 | 1 | 6767.99 | 6889.78 |
| **Ant.Cingulate - MFG** | | | | | | | |
| *PPC* | 1.19 | 1.08 | 1.34 | 0.05 | 1 | 7026.03 | 6288.96 |
| *PLI* | 1.09 | 0.95 | 1.28 | 0.08 | 1 | 5902.45 | 5968.35 |
| *IMCOH* | 1.08 | 0.99 | 1.22 | 0.05 | 1 | 6800.81 | 6500.42 |
| **OFC - Amygdala** | | | | | | | |
| *PPC* | 0.56 | 0.45 | 0.69 | 0.11 | 1 | 5318.97 | 7254.94 |
| *PLI* | 0.69 | 0.54 | 0.85 | 0.11 | 1 | 6212.42 | 6675.31 |
| *IMCOH* | 0.81 | 0.69 | 0.94 | 0.08 | 1 | 8111.63 | 7725.18 |
| **OFC - MFG** | | | | | | | |
| *PPC* | 1.30 | 1.10 | 1.51 | 0.08 | 1 | 3676.91 | 4978.70 |
| *PLI* | 1.19 | 1.00 | 1.41 | 0.09 | 1 | 4689.38 | 6029.14 |
| *IMCOH* | 1.14 | 1.03 | 1.27 | 0.05 | 1 | 5719.22 | 7167.14 |
| **Amygdala - MFG** | | | | | | | |
| *PPC* | 2.31 | 1.92 | 2.85 | 0.10 | 1 | 5016.48 | 6243.68 |
| *PLI* | 1.72 | 1.42 | 2.17 | 0.11 | 1 | 4595.40 | 5458.90 |
| *IMCOH* | 1.41 | 1.23 | 1.65 | 0.07 | 1 | 5789.61 | 6548.59 |

**Random effect of subject (20 levels)**

| Metric | Estimate | Est.Error | l-95% CI | u-95% CI | Rhat | Bulk_ESS | Tail_ESS |
|---|---|---|---|---|---|---|---|
| sd(Intercept) | | | | | | | |
| PPC | 0.29 | 0.07 | 0.18 | 0.45 | 1 | 3748.35 | 5679.98 |
| PLI | 0.24 | 0.07 | 0.12 | 0.39 | 1 | 4017.02 | 5564.17 |
| IMCOH | 0.12 | 0.04 | 0.05 | 0.22 | 1 | 3357.04 | 4429.69 |
| sd(OFC) | | | | | | | |
| PPC | 0.25 | 0.08 | 0.11 | 0.44 | 1 | 3628.50 | 4437.37 |
| PLI | 0.29 | 0.08 | 0.16 | 0.47 | 1 | 4369.55 | 5482.62 |
| IMCOH | 0.15 | 0.06 | 0.04 | 0.28 | 1 | 2799.17 | 2307.15 |
| sd(Ant.Cingulate) | | | | | | | |
| PPC | 0.08 | 0.06 | 0.00 | 0.24 | 1 | 3797.27 | 5666.12 |
| PLI | 0.17 | 0.09 | 0.02 | 0.39 | 1 | 2815.46 | 3806.63 |
| IMCOH | 0.11 | 0.06 | 0.01 | 0.24 | 1 | 2954.09 | 4096.15 |
| sd(Amygdala) | | | | | | | |
| PPC | 0.20 | 0.14 | 0.01 | 0.52 | 1 | 2086.02 | 4614.52 |
| PLI | 0.31 | 0.15 | 0.04 | 0.64 | 1 | 1966.57 | 2511.74 |
| IMCOH | 0.21 | 0.10 | 0.03 | 0.44 | 1 | 1813.88 | 2374.57 |
| cor(Intercept,OFC) | | | | | | | |
| PPC | 0.13 | 0.26 | -0.39 | 0.59 | 1 | 5251.32 | 7080.77 |
| PLI | -0.02 | 0.28 | -0.54 | 0.51 | 1 | 5024.44 | 7171.87 |
| IMCOH | 0.02 | 0.31 | -0.57 | 0.60 | 1 | 5912.54 | 6861.61 |
| cor(Intercept,Ant.Cingulate) | | | | | | | |
| PPC | 0.14 | 0.35 | -0.56 | 0.76 | 1 | 13034.49 | 8270.30 |
| PLI | -0.07 | 0.32 | -0.65 | 0.55 | 1 | 9686.27 | 8163.93 |
| IMCOH | -0.15 | 0.33 | -0.74 | 0.53 | 1 | 9040.82 | 8588.26 |
| cor(OFC,Ant.Cingulate) | | | | | | | |
| PPC | 0.11 | 0.38 | -0.64 | 0.77 | 1 | 9704.13 | 8504.71 |
| PLI | 0.18 | 0.34 | -0.51 | 0.76 | 1 | 8483.62 | 8719.18 |
| IMCOH | 0.27 | 0.35 | -0.47 | 0.85 | 1 | 8258.09 | 8935.83 |
| cor(Intercept,Amygdala) | | | | | | | |
| PPC | 0.17 | 0.33 | -0.51 | 0.75 | 1 | 8079.73 | 7855.12 |
| PLI | 0.22 | 0.32 | -0.47 | 0.77 | 1 | 6362.51 | 7114.38 |
| IMCOH | 0.15 | 0.35 | -0.55 | 0.76 | 1 | 5581.50 | 6575.69 |
| cor(OFC,Amygdala) | | | | | | | |
| PPC | 0.03 | 0.35 | -0.65 | 0.67 | 1 | 6815.07 | 8189.20 |
| PLI | 0.31 | 0.29 | -0.33 | 0.78 | 1 | 6342.14 | 7461.43 |
| IMCOH | 0.28 | 0.32 | -0.40 | 0.79 | 1 | 5504.73 | 8112.79 |
| cor(Ant.Cingulate,Amygdala) | | | | | | | |
| PPC | 0.05 | 0.37 | -0.67 | 0.72 | 1 | 5151.25 | 7900.87 |
| PLI | 0.12 | 0.34 | -0.55 | 0.74 | 1 | 5383.14 | 7518.80 |
| IMCOH | 0.14 | 0.35 | -0.57 | 0.77 | 1 | 5780.17 | 8613.64 |

##### OFC Coherence

| Metric | Mean | l-95% CI | u-95% CI | Est.Error | Rhat | Bulk_ESS | Tail_ESS |
|---|---|---|---|---|---|---|---|
| **Hippocampus - Ant.Cingulate** | | | | | | | |
| *PPC* | 0.60 | 0.44 | 0.81 | 0.15 | 1 | 5702.87 | 6957.51 |
| *PLI* | 0.71 | 0.54 | 0.93 | 0.14 | 1 | 4627.79 | 6553.62 |
| *IMCOH* | 0.76 | 0.64 | 0.90 | 0.09 | 1 | 5733.13 | 7804.48 |
| **Hippocampus - Amygdala** | | | | | | | |
| *PPC* | 0.84 | 0.62 | 1.14 | 0.16 | 1 | 4270.48 | 3615.85 |
| *PLI* | 0.96 | 0.74 | 1.23 | 0.13 | 1 | 5380.69 | 6934.09 |
| *IMCOH* | 0.98 | 0.84 | 1.14 | 0.08 | 1 | 7321.22 | 8125.72 |
| **Hippocampus - MFG** | | | | | | | |
| *PPC* | 0.83 | 0.64 | 1.09 | 0.13 | 1 | 4204.27 | 5925.53 |
| *PLI* | 0.92 | 0.70 | 1.19 | 0.13 | 1 | 3343.17 | 5727.74 |
| *IMCOH* | 0.92 | 0.79 | 1.07 | 0.08 | 1 | 3571.11 | 6210.78 |
| **Ant.Cingulate - Amygdala** | | | | | | | |
| *PPC* | 1.40 | 1.05 | 1.91 | 0.15 | 1 | 5596.56 | 4745.31 |
| *PLI* | 1.34 | 1.04 | 1.77 | 0.14 | 1 | 4393.30 | 5816.10 |
| *IMCOH* | 1.28 | 1.09 | 1.50 | 0.08 | 1 | 4920.29 | 6442.41 |
| **Ant.Cingulate - MFG** | | | | | | | |
| *PPC* | 1.39 | 1.12 | 1.77 | 0.12 | 1 | 4628.01 | 6513.77 |
| *PLI* | 1.29 | 1.07 | 1.56 | 0.09 | 1 | 4753.71 | 6422.57 |
| *IMCOH* | 1.20 | 1.07 | 1.36 | 0.06 | 1 | 5742.93 | 7189.46 |
| **Amygdala - MFG** | | | | | | | |
| *PPC* | 0.99 | 0.77 | 1.27 | 0.13 | 1 | 3307.70 | 2230.93 |
| *PLI* | 0.96 | 0.73 | 1.23 | 0.13 | 1 | 3119.64 | 5828.40 |
| *IMCOH* | 0.94 | 0.82 | 1.07 | 0.07 | 1 | 2649.24 | 5741.64 |

**Random effect of subject (20 levels)**

| Metric | Estimate | Est.Error | l-95% CI | u-95% CI | Rhat | Bulk_ESS | Tail_ESS |
|---|---|---|---|---|---|---|---|
| sd(Intercept) | | | | | | | |
| PPC | 0.35 | 0.08 | 0.22 | 0.52 | 1.00 | 3688.65 | 6006.78 |
| PLI | 0.31 | 0.07 | 0.19 | 0.46 | 1.00 | 3031.44 | 4867.58 |
| IMCOH | 0.16 | 0.04 | 0.09 | 0.26 | 1.00 | 3317.02 | 5501.44 |
| sd(Amygdala) | | | | | | | |
| PPC | 0.22 | 0.16 | 0.01 | 0.57 | 1.00 | 1868.88 | 2442.13 |
| PLI | 0.41 | 0.13 | 0.17 | 0.68 | 1.00 | 1410.72 | 1928.41 |
| IMCOH | 0.18 | 0.08 | 0.03 | 0.35 | 1.01 | 767.60 | 1921.54 |
| sd(Ant.Cingulate) | | | | | | | |
| PPC | 0.31 | 0.13 | 0.10 | 0.60 | 1.00 | 2833.34 | 2919.60 |
| PLI | 0.27 | 0.09 | 0.13 | 0.49 | 1.00 | 4206.18 | 5717.90 |
| IMCOH | 0.16 | 0.06 | 0.07 | 0.32 | 1.00 | 4769.29 | 6105.75 |
| sd(Hippocampus) | | | | | | | |
| PPC | 0.46 | 0.12 | 0.26 | 0.72 | 1.00 | 3952.65 | 6351.04 |
| PLI | 0.47 | 0.11 | 0.29 | 0.71 | 1.00 | 2554.97 | 4517.03 |
| IMCOH | 0.26 | 0.08 | 0.13 | 0.43 | 1.00 | 1963.87 | 4274.65 |
| cor(Intercept,Amygdala) | | | | | | | |
| PPC | -0.04 | 0.34 | -0.66 | 0.63 | 1.00 | 7902.29 | 7645.63 |
| PLI | -0.23 | 0.26 | -0.68 | 0.34 | 1.00 | 2371.48 | 4767.25 |
| IMCOH | -0.22 | 0.31 | -0.73 | 0.45 | 1.00 | 2070.52 | 5501.62 |
| cor(Intercept,Ant.Cingulate) | | | | | | | |
| PPC | -0.19 | 0.29 | -0.71 | 0.39 | 1.00 | 5848.65 | 7346.47 |
| PLI | -0.28 | 0.26 | -0.72 | 0.27 | 1.00 | 5145.33 | 7496.02 |
| IMCOH | -0.24 | 0.28 | -0.74 | 0.36 | 1.00 | 5812.89 | 7629.37 |
| cor(Amygdala,Ant.Cingulate) | | | | | | | |
| PPC | 0.09 | 0.37 | -0.63 | 0.74 | 1.00 | 3055.61 | 5312.81 |
| PLI | 0.44 | 0.31 | -0.29 | 0.90 | 1.00 | 3528.05 | 5921.26 |
| IMCOH | 0.23 | 0.38 | -0.55 | 0.84 | 1.00 | 3294.74 | 6709.52 |
| cor(Intercept,Hippocampus) | | | | | | | |
| PPC | -0.35 | 0.23 | -0.72 | 0.14 | 1.00 | 4838.11 | 7237.37 |
| PLI | -0.56 | 0.19 | -0.85 | -0.12 | 1.00 | 2882.36 | 5046.87 |
| IMCOH | -0.52 | 0.23 | -0.86 | 0.01 | 1.00 | 2784.98 | 5784.82 |
| cor(Amygdala,Hippocampus) | | | | | | | |
| PPC | 0.09 | 0.35 | -0.60 | 0.71 | 1.00 | 2329.42 | 4936.75 |
| PLI | 0.57 | 0.21 | 0.07 | 0.89 | 1.00 | 3307.32 | 5101.33 |
| IMCOH | 0.49 | 0.27 | -0.15 | 0.88 | 1.00 | 2502.75 | 3280.00 |
| cor(Ant.Cingulate,Hippocampus) | | | | | | | |
| PPC | 0.22 | 0.30 | -0.40 | 0.73 | 1.00 | 3548.29 | 6451.91 |
| PLI | 0.33 | 0.27 | -0.25 | 0.77 | 1.00 | 4301.43 | 6704.44 |
| IMCOH | 0.24 | 0.31 | -0.41 | 0.76 | 1.00 | 4206.40 | 6833.76 |

##### ACC Coherence

| Metric | Mean | l-95% CI | u-95% CI | Est.Error | Rhat | Bulk_ESS | Tail_ESS |
|---|---|---|---|---|---|---|---|
| **Hippocampus - OFC** | | | | | | | |
| *PPC* | 0.54 | 0.40 | 0.74 | 0.16 | 1 | 5344.18 | 7070.50 |
| *PLI* | 0.69 | 0.53 | 0.91 | 0.14 | 1 | 6929.50 | 8427.96 |
| *IMCOH* | 0.76 | 0.65 | 0.91 | 0.08 | 1 | 5089.88 | 6720.10 |
| **Hippocampus - Amygdala** | | | | | | | |
| *PPC* | 0.95 | 0.69 | 1.33 | 0.17 | 1 | 7274.48 | 6570.81 |
| *PLI* | 1.00 | 0.73 | 1.40 | 0.16 | 1 | 7591.34 | 8566.53 |
| *IMCOH* | 1.02 | 0.85 | 1.22 | 0.09 | 1 | 5739.08 | 7120.92 |
| **Hippocampus - MFG** | | | | | | | |
| *PPC* | 0.56 | 0.40 | 0.80 | 0.17 | 1 | 3424.54 | 5558.90 |
| *PLI* | 0.69 | 0.52 | 0.94 | 0.15 | 1 | 4415.56 | 6450.35 |
| *IMCOH* | 0.77 | 0.65 | 0.91 | 0.09 | 1 | 3670.91 | 5033.13 |
| **OFC - Amygdala** | | | | | | | |
| *PPC* | 1.75 | 1.25 | 2.44 | 0.17 | 1 | 7180.89 | 7478.92 |
| *PLI* | 1.45 | 1.11 | 1.87 | 0.13 | 1 | 7740.18 | 8798.56 |
| *IMCOH* | 1.33 | 1.15 | 1.54 | 0.07 | 1 | 7833.11 | 7786.94 |
| **OFC - MFG** | | | | | | | |
| *PPC* | 1.02 | 0.78 | 1.40 | 0.15 | 1 | 2893.81 | 4815.41 |
| *PLI* | 1.00 | 0.82 | 1.25 | 0.11 | 1 | 4044.41 | 6261.57 |
| *IMCOH* | 1.01 | 0.90 | 1.13 | 0.06 | 1 | 4296.64 | 3799.16 |
| **Amygdala - MFG** | | | | | | | |
| *PPC* | 0.58 | 0.43 | 0.83 | 0.17 | 1 | 4037.23 | 6537.76 |
| *PLI* | 0.69 | 0.53 | 0.90 | 0.13 | 1 | 5225.58 | 6546.13 |
| *IMCOH* | 0.76 | 0.65 | 0.88 | 0.07 | 1 | 4476.46 | 5066.12 |

**Random effect of subject (20 levels)**

| Metric | Estimate | Est.Error | l-95% CI | u-95% CI | Rhat | Bulk_ESS | Tail_ESS |
|---|---|---|---|---|---|---|---|
| sd(Intercept) | | | | | | | |
| PPC | 0.36 | 0.09 | 0.21 | 0.56 | 1 | 3052.36 | 5043.26 |
| PLI | 0.29 | 0.08 | 0.17 | 0.47 | 1 | 3258.38 | 4776.32 |
| IMCOH | 0.14 | 0.04 | 0.07 | 0.23 | 1 | 2610.22 | 3246.29 |
| sd(OFC) | | | | | | | |
| PPC | 0.32 | 0.15 | 0.05 | 0.66 | 1 | 1484.84 | 1879.30 |
| PLI | 0.17 | 0.12 | 0.01 | 0.43 | 1 | 1810.93 | 3474.33 |
| IMCOH | 0.07 | 0.06 | 0.00 | 0.22 | 1 | 2353.85 | 3539.85 |
| sd(Hippocampus) | | | | | | | |
| PPC | 0.47 | 0.13 | 0.26 | 0.76 | 1 | 2597.73 | 3738.67 |
| PLI | 0.42 | 0.12 | 0.22 | 0.68 | 1 | 3009.12 | 5013.76 |
| IMCOH | 0.21 | 0.07 | 0.08 | 0.38 | 1 | 3124.62 | 4092.28 |
| sd(Amygdala) | | | | | | | |
| PPC | 0.36 | 0.17 | 0.04 | 0.72 | 1 | 1889.57 | 2078.11 |
| PLI | 0.21 | 0.15 | 0.01 | 0.56 | 1 | 2397.63 | 5220.18 |
| IMCOH | 0.08 | 0.07 | 0.00 | 0.25 | 1 | 3007.89 | 4035.15 |
| cor(Intercept,OFC) | | | | | | | |
| PPC | -0.28 | 0.30 | -0.76 | 0.37 | 1 | 3971.41 | 5153.54 |
| PLI | -0.17 | 0.34 | -0.74 | 0.55 | 1 | 5786.63 | 7496.21 |
| IMCOH | -0.08 | 0.37 | -0.75 | 0.64 | 1 | 2703.22 | 3110.21 |
| cor(Intercept,Hippocampus) | | | | | | | |
| PPC | -0.47 | 0.24 | -0.83 | 0.08 | 1 | 3661.74 | 5458.28 |
| PLI | -0.57 | 0.23 | -0.89 | -0.03 | 1 | 4434.19 | 7086.97 |
| IMCOH | -0.51 | 0.26 | -0.88 | 0.08 | 1 | 3623.48 | 6049.97 |
| cor(OFC,Hippocampus) | | | | | | | |
| PPC | 0.43 | 0.29 | -0.23 | 0.87 | 1 | 3037.11 | 4375.53 |
| PLI | 0.32 | 0.34 | -0.46 | 0.83 | 1 | 2910.66 | 4921.02 |
| IMCOH | 0.06 | 0.36 | -0.65 | 0.73 | 1 | 4170.58 | 4719.64 |
| cor(Intercept,Amygdala) | | | | | | | |
| PPC | -0.27 | 0.30 | -0.76 | 0.38 | 1 | 4766.07 | 6322.43 |
| PLI | -0.18 | 0.34 | -0.75 | 0.54 | 1 | 5896.21 | 7785.74 |
| IMCOH | -0.10 | 0.37 | -0.74 | 0.64 | 1 | 4131.98 | 2646.67 |
| cor(OFC,Amygdala) | | | | | | | |
| PPC | 0.32 | 0.33 | -0.40 | 0.85 | 1 | 3948.45 | 5758.67 |
| PLI | 0.20 | 0.39 | -0.60 | 0.82 | 1 | 4375.03 | 8330.55 |
| IMCOH | 0.08 | 0.39 | -0.69 | 0.76 | 1 | 2431.99 | 3113.18 |
| cor(Hippocampus,Amygdala) | | | | | | | |
| PPC | 0.53 | 0.28 | -0.17 | 0.91 | 1 | 3836.34 | 4372.18 |
| PLI | 0.25 | 0.35 | -0.52 | 0.81 | 1 | 5700.42 | 8244.21 |
| IMCOH | 0.14 | 0.37 | -0.61 | 0.78 | 1 | 5589.29 | 5796.30 |

##### MFG Coherence

| Metric | Mean | l-95% CI | u-95% CI | Est.Error | Rhat | Bulk_ESS | Tail_ESS |
|---|---|---|---|---|---|---|---|
| **Hippocampus - Ant.Cingulate** | | | | | | | |
| *PPC* | 0.48 | 0.33 | 0.69 | 0.18 | 1 | 4195.15 | 6064.65 |
| *PLI* | 0.60 | 0.44 | 0.83 | 0.16 | 1 | 5394.40 | 6742.99 |
| *IMCOH* | 0.69 | 0.57 | 0.83 | 0.09 | 1 | 7989.21 | 7723.94 |
| **Hippocampus - Amygdala** | | | | | | | |
| *PPC* | 0.96 | 0.79 | 1.18 | 0.10 | 1 | 6659.15 | 7224.91 |
| *PLI* | 1.04 | 0.85 | 1.29 | 0.11 | 1 | 8904.23 | 7898.83 |
| *IMCOH* | 1.03 | 0.90 | 1.18 | 0.07 | 1 | 10835.17 | 8653.30 |
| **Hippocampus - OFC** | | | | | | | |
| *PPC* | 0.59 | 0.49 | 0.71 | 0.09 | 1 | 3791.46 | 3680.20 |
| *PLI* | 0.76 | 0.63 | 0.93 | 0.10 | 1 | 4617.98 | 5620.85 |
| *IMCOH* | 0.80 | 0.72 | 0.89 | 0.06 | 1 | 7980.87 | 7758.26 |
| **Ant.Cingulate - Amygdala** | | | | | | | |
| *PPC* | 2.02 | 1.41 | 2.87 | 0.18 | 1 | 3489.54 | 6514.98 |
| *PLI* | 1.73 | 1.27 | 2.35 | 0.16 | 1 | 5950.72 | 7320.89 |
| *IMCOH* | 1.50 | 1.23 | 1.83 | 0.10 | 1 | 8056.84 | 7656.85 |
| **Ant.Cingulate - OFC** | | | | | | | |
| *PPC* | 1.24 | 0.89 | 1.70 | 0.16 | 1 | 3603.86 | 6400.16 |
| *PLI* | 1.27 | 0.98 | 1.63 | 0.13 | 1 | 5928.85 | 6999.18 |
| *IMCOH* | 1.17 | 1.00 | 1.35 | 0.08 | 1 | 7590.22 | 5705.05 |
| **Amygdala - OFC** | | | | | | | |
| *PPC* | 0.61 | 0.51 | 0.74 | 0.09 | 1 | 4508.78 | 6163.37 |
| *PLI* | 0.73 | 0.60 | 0.89 | 0.10 | 1 | 6344.24 | 6881.44 |
| *IMCOH* | 0.78 | 0.68 | 0.89 | 0.07 | 1 | 8278.89 | 7470.18 |

**Random effect of subject (20 levels)**

| Metric | Estimate | Est.Error | l-95% CI | u-95% CI | Rhat | Bulk_ESS | Tail_ESS |
|---|---|---|---|---|---|---|---|
| sd(Intercept) | | | | | | | |
| PPC | 0.29 | 0.07 | 0.18 | 0.45 | 1 | 3709.08 | 6398.23 |
| PLI | 0.27 | 0.06 | 0.17 | 0.41 | 1 | 4107.75 | 6264.62 |
| IMCOH | 0.14 | 0.04 | 0.08 | 0.23 | 1 | 4147.85 | 6502.18 |
| sd(Ant.Cingulate) | | | | | | | |
| PPC | 0.45 | 0.15 | 0.21 | 0.81 | 1 | 3579.58 | 4839.25 |
| PLI | 0.31 | 0.14 | 0.05 | 0.63 | 1 | 2673.88 | 2214.04 |
| IMCOH | 0.18 | 0.09 | 0.02 | 0.38 | 1 | 3228.28 | 3460.52 |
| sd(Hippocampus) | | | | | | | |
| PPC | 0.25 | 0.09 | 0.09 | 0.45 | 1 | 2790.10 | 3890.98 |
| PLI | 0.30 | 0.09 | 0.14 | 0.51 | 1 | 3051.99 | 4867.98 |
| IMCOH | 0.14 | 0.06 | 0.03 | 0.28 | 1 | 2689.03 | 2931.43 |
| sd(Amygdala) | | | | | | | |
| PPC | 0.18 | 0.12 | 0.01 | 0.45 | 1 | 1292.58 | 4314.75 |
| PLI | 0.23 | 0.12 | 0.02 | 0.47 | 1 | 2319.28 | 3181.68 |
| IMCOH | 0.18 | 0.08 | 0.02 | 0.34 | 1 | 2141.11 | 2272.28 |
| cor(Intercept,Ant.Cingulate) | | | | | | | |
| PPC | -0.11 | 0.27 | -0.59 | 0.43 | 1 | 4961.26 | 7723.10 |
| PLI | 0.02 | 0.29 | -0.54 | 0.58 | 1 | 7964.54 | 8010.70 |
| IMCOH | 0.00 | 0.31 | -0.59 | 0.61 | 1 | 8280.11 | 5775.10 |
| cor(Intercept,Hippocampus) | | | | | | | |
| PPC | -0.40 | 0.25 | -0.78 | 0.18 | 1 | 5123.57 | 6798.39 |
| PLI | -0.54 | 0.21 | -0.86 | -0.05 | 1 | 5814.22 | 7747.89 |
| IMCOH | -0.45 | 0.28 | -0.86 | 0.20 | 1 | 5359.10 | 7534.82 |
| cor(Ant.Cingulate,Hippocampus) | | | | | | | |
| PPC | -0.20 | 0.31 | -0.75 | 0.40 | 1 | 4440.56 | 7665.34 |
| PLI | -0.25 | 0.30 | -0.77 | 0.36 | 1 | 4882.01 | 6109.28 |
| IMCOH | -0.26 | 0.32 | -0.81 | 0.39 | 1 | 7044.30 | 6719.57 |
| cor(Intercept,Amygdala) | | | | | | | |
| PPC | -0.20 | 0.34 | -0.76 | 0.51 | 1 | 3503.79 | 7201.54 |
| PLI | -0.31 | 0.30 | -0.78 | 0.37 | 1 | 6119.03 | 5739.17 |
| IMCOH | -0.34 | 0.29 | -0.80 | 0.31 | 1 | 6349.54 | 7389.87 |
| cor(Ant.Cingulate,Amygdala) | | | | | | | |
| PPC | -0.02 | 0.36 | -0.68 | 0.65 | 1 | 1951.86 | 2959.99 |
| PLI | -0.16 | 0.36 | -0.78 | 0.56 | 1 | 5661.35 | 6870.25 |
| IMCOH | -0.16 | 0.35 | -0.78 | 0.55 | 1 | 6674.35 | 5816.78 |
| cor(Hippocampus,Amygdala) | | | | | | | |
| PPC | 0.24 | 0.35 | -0.50 | 0.80 | 1 | 4727.17 | 7692.15 |
| PLI | 0.45 | 0.30 | -0.28 | 0.88 | 1 | 5197.40 | 6705.37 |
| IMCOH | 0.37 | 0.31 | -0.33 | 0.86 | 1 | 5574.81 | 6613.42 |

**Supp. Table 3.** Posterior predictive checks for regional theta coherence analyses

| Posterior Predictive Check: Amygdala | | | | |
|---|---|---|---|---|
| Statistic | Observed Value | Posterior Predictive Mean | Credible Interval | Posterior Predictive p value |
| **PPC** | | | | |
| *Mean* | -5.0112562 | -5.0110895 | [-5.044, -4.976] | 0.52 |
| *Standard Deviation* | 0.9911363 | 0.9943510 | [0.963, 1.029] | 0.55 |
| **PLI** | | | | |
| *Mean* | -3.9392873 | -3.9370651 | [-3.97, -3.9] | 0.54 |
| *Standard Deviation* | 0.8322597 | 0.8345424 | [0.803, 0.868] | 0.61 |
| **IMCOH** | | | | |
| *Mean* | -3.0220568 | -3.0214119 | [-3.046, -2.999] | 0.56 |
| *Standard Deviation* | 0.5608367 | 0.5615798 | [0.544, 0.583] | 0.51 |

| Posterior Predictive Check: Hippocampus | | | | |
|---|---|---|---|---|
| Statistic | Observed Value | Posterior Predictive Mean | Credible Interval | Posterior Predictive p value |
| **PPC** | | | | |
| *Mean* | -5.0983353 | -5.0982825 | [-5.125, -5.07] | 0.49 |
| *Standard Deviation* | 0.8094896 | 0.8110096 | [0.786, 0.833] | 0.57 |
| **PLI** | | | | |
| *Mean* | -3.9392681 | -3.9392302 | [-3.968, -3.911] | 0.49 |
| *Standard Deviation* | 0.6875272 | 0.6884490 | [0.667, 0.709] | 0.52 |
| **IMCOH** | | | | |
| *Mean* | -3.0305881 | -3.0294654 | [-3.054, -3.006] | 0.53 |
| *Standard Deviation* | 0.4844154 | 0.4868016 | [0.47, 0.502] | 0.63 |

| Posterior Predictive Check: OFC | | | | |
|---|---|---|---|---|
| Statistic | Observed Value | Posterior Predictive Mean | Credible Interval | Posterior Predictive p value |
| **PPC** | | | | |
| *Mean* | -4.9325216 | -4.9332967 | [-4.958, -4.906] | 0.47 |
| *Standard Deviation* | 0.8963513 | 0.8985080 | [0.877, 0.923] | 0.58 |
| **PLI** | | | | |
| *Mean* | -3.8465322 | -3.8469478 | [-3.873, -3.823] | 0.47 |
| *Standard Deviation* | 0.7881697 | 0.7915835 | [0.77, 0.812] | 0.62 |
| **IMCOH** | | | | |
| *Mean* | -2.9533249 | -2.9525753 | [-2.972, -2.934] | 0.55 |
| *Standard Deviation* | 0.5282490 | 0.5298779 | [0.515, 0.548] | 0.58 |

| Posterior Predictive Check: ACC | | | | |
|---|---|---|---|---|
| Statistic | Observed Value | Posterior Predictive Mean | Credible Interval | Posterior Predictive p value |
| **PPC** | | | | |
| *Mean* | -4.7461005 | -4.7489152 | [-4.796, -4.698] | 0.45 |
| *Standard Deviation* | 0.9938289 | 0.9943672 | [0.963, 1.029] | 0.51 |
| **PLI** | | | | |
| *Mean* | -3.6963435 | -3.6954031 | [-3.739, -3.65] | 0.49 |
| *Standard Deviation* | 0.9024046 | 0.9050765 | [0.874, 0.935] | 0.58 |
| **IMCOH** | | | | |
| *Mean* | -2.8681809 | -2.8702762 | [-2.899, -2.845] | 0.48 |
| *Standard Deviation* | 0.5755032 | 0.5770774 | [0.553, 0.601] | 0.53 |

| Posterior Predictive Check: MFG | | | | |
|---|---|---|---|---|
| Statistic | Observed Value | Posterior Predictive Mean | Credible Interval | Posterior Predictive p value |
| **PPC** | | | | |
| *Mean* | -4.9665244 | -4.9665264 | [-4.995, -4.936] | 0.49 |
| *Standard Deviation* | 0.9231406 | 0.9245712 | [0.898, 0.954] | 0.55 |
| **PLI** | | | | |
| *Mean* | -3.8542984 | -3.8575713 | [-3.883, -3.836] | 0.43 |
| *Standard Deviation* | 0.8024406 | 0.8039547 | [0.783, 0.827] | 0.53 |
| **IMCOH** | | | | |
| *Mean* | -2.9851885 | -2.9830539 | [-3.004, -2.964] | 0.57 |
| *Standard Deviation* | 0.5273133 | 0.5290527 | [0.507, 0.547] | 0.59 |

**Supp Table 4.** Estimates & posterior predictive checks for models estimating the effect of time on theta coherence, including alternate timepoints & alternate coherence metrics

**Effect of Time on Theta Coherence**

| | | | Model Estimates | | | | | | | Posterior Predictive Checks | | | | | | |
|---|---|---|---|---|---|---|---|---|---|---|---|---|---|---|---|---|
| Metric | Timepoint | Estimate | Est.Error | l-95% CI | u-95% CI | Rhat | Bulk_ESS | Tail_ESS | True Mean | True SD | Posterior Predictive Mean | CI- Mean | Posterior Predictive SD | CI- SD | Post. Pred. Mean p value | Post. Pred. SD p value |
| **Approach** | | | | | | | | | | | | | | | | |
| *Im. Coh.* | *1.5 (s)* | 0.06 | 0.04 | -0.01 | 0.14 | 1.00 | 445.27 | 1332.97 | 0 | 1 | 0.000 | [-0.008, 0.008] | 1.000 | [0.994, 1.008] | 0.43 | 0.53 |
| *Im. Coh.* | *1.6 (s)* | 0.08 | 0.03 | 0.01 | 0.14 | 1.00 | 3196.19 | 5592.32 | 0 | 1 | 0.000 | [-0.008, 0.008] | 1.000 | [0.994, 1.005] | 0.50 | 0.50 |
| *Im. Coh.* | *1.7 (s)* | 0.09 | 0.03 | 0.03 | 0.14 | 1.00 | 4131.51 | 6590.06 | 0 | 1 | 0.000 | [-0.007, 0.009] | 1.001 | [0.993, 1.009] | 0.49 | 0.55 |
| *Im. Coh.* | *1.8 (s)* | 0.09 | 0.03 | 0.03 | 0.14 | 1.00 | 4093.91 | 6303.93 | 0 | 1 | -0.001 | [-0.009, 0.011] | 1.000 | [0.993, 1.006] | 0.40 | 0.49 |
| *Im. Coh.* | *2 (s)* | 0.09 | 0.02 | 0.05 | 0.14 | 1.00 | 7680.42 | 9704.80 | 0 | 1 | 0.000 | [-0.009, 0.007] | 1.000 | [0.992, 1.006] | 0.52 | 0.58 |
| *PPC* | *2 (s)* | 0.07 | 0.02 | 0.03 | 0.10 | 1.00 | 9293.28 | 8836.05 | 0 | 1 | -0.001 | [-0.008, 0.006] | 1.001 | [0.995, 1.007] | 0.43 | 0.57 |
| *PLI* | *2 (s)* | 0.08 | 0.02 | 0.04 | 0.13 | 1.00 | 10046.50 | 10629.59 | 0 | 1 | 0.000 | [-0.005, 0.006] | 1.000 | [0.994, 1.004] | 0.46 | 0.50 |
| **Avoid** | | | | | | | | | | | | | | | | |
| *Im. Coh.* | *1.5 (s)* | -0.13 | 0.04 | -0.20 | -0.05 | 1.00 | 5387.01 | 6839.51 | 0 | 1 | 0.000 | [-0.012, 0.009] | 1.001 | [0.993, 1.008] | 0.48 | 0.56 |
| *Im. Coh.* | *1.6 (s)* | -0.13 | 0.03 | -0.19 | -0.06 | 1.01 | 1080.11 | 2326.39 | 0 | 1 | 0.000 | [-0.009, 0.01] | 1.000 | [0.993, 1.008] | 0.55 | 0.51 |
| *Im. Coh.* | *1.7 (s)* | -0.12 | 0.03 | -0.18 | -0.06 | 1.00 | 4770.02 | 6872.33 | 0 | 1 | 0.000 | [-0.009, 0.007] | 1.000 | [0.993, 1.008] | 0.50 | 0.48 |
| *Im. Coh.* | *1.8 (s)* | -0.12 | 0.03 | -0.18 | -0.06 | 1.00 | 5651.34 | 7151.48 | 0 | 1 | 0.001 | [-0.006, 0.008] | 1.000 | [0.994, 1.007] | 0.56 | 0.42 |
| *Im. Coh.* | *2 (s)* | -0.11 | 0.03 | -0.16 | -0.06 | 1.00 | 3520.35 | 4769.50 | 0 | 1 | 0.000 | [-0.009, 0.009] | 1.000 | [0.993, 1.006] | 0.44 | 0.54 |
| *PPC* | *2 (s)* | -0.12 | 0.03 | -0.17 | -0.07 | 1.00 | 7365.40 | 10049.77 | 0 | 1 | 0.000 | [-0.005, 0.006] | 1.000 | [0.995, 1.005] | 0.49 | 0.51 |
| *PLI* | *2 (s)* | -0.11 | 0.03 | -0.18 | -0.05 | 1.00 | 5228.66 | 8321.24 | 0 | 1 | 0.000 | [-0.007, 0.007] | 1.000 | [0.994, 1.007] | 0.52 | 0.45 |

**Supp Table 5.** Full results for a model estimating the effect of approach time on theta synchrony, with interactions for each region pair

**Random effect of subject (15 levels)**

| | Estimate | Est.Error | l-95% CI | u-95% CI | Rhat | Bulk_ESS | Tail_ESS |
|---|---|---|---|---|---|---|---|
| *sd(Intercept)* | 0.09 | 0.02 | 0.07 | 0.13 | 1 | 3094.13 | 4861.83 |
| *sd(Time)* | 0.02 | 0.01 | 0.01 | 0.04 | 1 | 5082.88 | 8310.22 |
| *cor(Intercept,Time)* | 0.32 | 0.22 | -0.14 | 0.70 | 1 | 6190.50 | 7782.18 |

**Random effect of electrode pair (3119 levels)**

| | Estimate | Est.Error | l-95% CI | u-95% CI | Rhat | Bulk_ESS | Tail_ESS |
|---|---|---|---|---|---|---|---|
| *sd(Intercept)* | 0.14 | 0.0 | 0.13 | 0.14 | 1 | 5224.90 | 8229.24 |
| *sd(Time)* | 0.01 | 0.0 | 0.00 | 0.02 | 1 | 3797.34 | 4138.36 |
| *cor(Intercept,Time)* | 0.60 | 0.2 | 0.19 | 0.93 | 1 | 7906.15 | 6541.53 |

**Theta Model Estimates**

| Variable | Estimate | Est.Error | l-95% CI | u-95% CI | Rhat | Bulk_ESS | Tail_ESS |
|---|---|---|---|---|---|---|---|
| *Intercept* | -0.10 | 0.03 | -0.15 | -0.04 | 1 | 1724.61 | 2994.63 |
| *time* | 0.07 | 0.01 | 0.04 | 0.09 | 1 | 2816.57 | 4873.93 |
| *amyg_mfg* | -0.08 | 0.03 | -0.14 | -0.03 | 1 | 2300.09 | 4603.84 |
| *hc_amyg* | 0.10 | 0.02 | 0.05 | 0.15 | 1 | 2295.42 | 3978.02 |
| *hc_cing* | 0.03 | 0.03 | -0.02 | 0.09 | 1 | 2509.18 | 4633.75 |
| *hc_mfg* | -0.03 | 0.03 | -0.08 | 0.02 | 1 | 2350.43 | 4116.79 |
| *mfg_cing* | 0.08 | 0.02 | 0.04 | 0.13 | 1 | 2131.18 | 4117.48 |
| *ofc_amyg* | 0.05 | 0.02 | 0.00 | 0.09 | 1 | 2150.91 | 3522.45 |
| *ofc_cing* | 0.10 | 0.02 | 0.05 | 0.14 | 1 | 2102.71 | 3999.21 |
| *ofc_hc* | 0.07 | 0.02 | 0.02 | 0.11 | 1 | 2116.21 | 3496.72 |
| *ofc_mfg* | 0.08 | 0.02 | 0.04 | 0.13 | 1 | 2066.83 | 3851.51 |
| *time:amyg_mfg* | -0.06 | 0.01 | -0.09 | -0.03 | 1 | 3736.22 | 7119.85 |
| *time:hc_amyg* | -0.02 | 0.01 | -0.04 | 0.01 | 1 | 3269.16 | 6212.89 |
| *time:hc_cing* | -0.04 | 0.02 | -0.07 | -0.01 | 1 | 4250.59 | 7265.66 |
| *time:hc_mfg* | -0.05 | 0.01 | -0.08 | -0.02 | 1 | 3254.37 | 6105.83 |
| *time:mfg_cing* | -0.04 | 0.01 | -0.06 | -0.01 | 1 | 2819.12 | 5389.10 |
| *time:ofc_amyg* | -0.02 | 0.01 | -0.05 | 0.01 | 1 | 2986.13 | 5495.49 |
| *time:ofc_cing* | -0.01 | 0.01 | -0.03 | 0.02 | 1 | 3029.68 | 5611.96 |
| *time:ofc_hc* | -0.03 | 0.01 | -0.06 | 0.00 | 1 | 2986.60 | 5425.94 |
| *time:ofc_mfg* | -0.01 | 0.01 | -0.04 | 0.01 | 1 | 2828.36 | 5684.88 |

**Supp Table 6.** Full results for a model estimating the effect of approach time on HFA synchrony, with interactions for each region pair

**Random effect of subject (15 levels)**

| | Estimate | Est.Error | l-95% CI | u-95% CI | Rhat | Bulk_ESS | Tail_ESS |
|---|---|---|---|---|---|---|---|
| *sd(Intercept)* | 0.05 | 0.01 | 0.03 | 0.08 | 1 | 2921.06 | 5900.38 |
| *sd(Time)* | 0.01 | 0.00 | 0.00 | 0.02 | 1 | 4493.44 | 5419.55 |
| *cor(Intercept,Time)* | 0.33 | 0.31 | -0.31 | 0.86 | 1 | 8858.24 | 7213.78 |

**Random effect of electrode pair (3119 levels)**

| | Estimate | Est.Error | l-95% CI | u-95% CI | Rhat | Bulk_ESS | Tail_ESS |
|---|---|---|---|---|---|---|---|
| *sd(Intercept)* | 0.09 | 0.00 | 0.09 | 0.10 | 1 | 6453.04 | 8608.55 |
| *sd(Time)* | 0.02 | 0.00 | 0.01 | 0.03 | 1 | 3458.02 | 6369.21 |
| *cor(Intercept,Time)* | -0.81 | 0.09 | -0.96 | -0.61 | 1 | 3029.59 | 5184.90 |

**HFA Model Estimates**

| Variable | Estimate | Est.Error | l-95% CI | u-95% CI | Rhat | Bulk_ESS | Tail_ESS |
|---|---|---|---|---|---|---|---|
| *Intercept* | 0.00 | 0.02 | -0.04 | 0.04 | 1 | 2163.22 | 4013.68 |
| *time* | -0.01 | 0.01 | -0.04 | 0.01 | 1 | 2159.44 | 4148.57 |
| *amyg_mfg* | -0.04 | 0.02 | -0.09 | 0.00 | 1 | 2634.48 | 4879.21 |
| *hc_amyg* | 0.11 | 0.02 | 0.07 | 0.14 | 1 | 2316.04 | 4735.18 |
| *hc_cing* | -0.01 | 0.02 | -0.05 | 0.03 | 1 | 2594.82 | 5928.56 |
| *hc_mfg* | -0.05 | 0.02 | -0.09 | -0.01 | 1 | 2365.76 | 5141.01 |
| *mfg_cing* | 0.00 | 0.02 | -0.04 | 0.03 | 1 | 2175.58 | 4353.26 |
| *ofc_amyg* | 0.03 | 0.02 | 0.00 | 0.07 | 1 | 2270.31 | 4293.34 |
| *ofc_cing* | 0.00 | 0.02 | -0.04 | 0.03 | 1 | 2152.27 | 5120.89 |
| *ofc_hc* | 0.02 | 0.02 | -0.01 | 0.06 | 1 | 2170.50 | 4771.69 |
| *ofc_mfg* | 0.02 | 0.02 | -0.02 | 0.05 | 1 | 2002.56 | 4645.52 |
| *time:amyg_mfg* | 0.03 | 0.01 | 0.00 | 0.06 | 1 | 3023.05 | 5478.40 |
| *time:hc_amyg* | 0.02 | 0.01 | 0.00 | 0.05 | 1 | 2420.92 | 4612.21 |
| *time:hc_cing* | 0.02 | 0.02 | -0.01 | 0.05 | 1 | 3543.39 | 6615.43 |
| *time:hc_mfg* | -0.01 | 0.01 | -0.03 | 0.02 | 1 | 2633.62 | 4879.35 |
| *time:mfg_cing* | 0.02 | 0.01 | 0.00 | 0.04 | 1 | 2256.33 | 4578.76 |
| *time:ofc_amyg* | 0.01 | 0.01 | -0.01 | 0.04 | 1 | 2482.32 | 4461.08 |
| *time:ofc_cing* | 0.03 | 0.01 | 0.01 | 0.06 | 1 | 2269.69 | 4619.77 |
| *time:ofc_hc* | 0.02 | 0.01 | 0.00 | 0.04 | 1 | 2385.92 | 4812.98 |
| *time:ofc_mfg* | 0.03 | 0.01 | 0.00 | 0.05 | 1 | 2209.27 | 4430.00 |

**Supp Table 7.** Posterior predictive checks for models estimating the effect of approach time on HFA & Theta synchrony

**Posterior Predictive Check: Synchrony ~ Approach Time**

| Statistic | Observed Value | Posterior Predictive Mean | Credible Interval | Posterior Predictive p value |
|---|---|---|---|---|
| **Theta** | | | | |
| *Mean* | 0.01 | 0.01 | [0.007, 0.017] | 0.45 |
| *Standard Deviation* | 1.00 | 1.00 | [1.002, 1.009] | 0.45 |
| **HFA** | | | | |
| *Mean* | 0.02 | 0.02 | [0.011, 0.02] | 0.51 |
| *Standard Deviation* | 0.99 | 0.99 | [0.982, 0.988] | 0.51 |
